# Voltage imaging in *Drosophila* using a hybrid chemical-genetic rhodamine voltage reporter

**DOI:** 10.1101/2021.08.05.455279

**Authors:** Molly J. Kirk, Brittany R. Benlian, Yifu Han, Arya Gold, Ashvin Ravi, Parker E. Deal, Rosana S. Molina, Mikhail Drobizhev, Dion Dickman, Kristin Scott, Evan W. Miller

**Author notes:** **Contact Info:** email addresses. **Author Contributions:** Designed experiments – MJK, BRB, YH, MD, DD, KS, EWM. Performed Experiments – MJK, BRB, YH, AR, AG, RSM, MD. Contributed Unpublished Reagents/Analysis – PED, MD, MJK, BRB. Analyzed Data – MJK, BRB, YH, RSM, MD, DD, KS, EWM. Wrote Manuscript – MJK, BRB, YH, DD, KS, EWM.

## Abstract

We combine a chemically-synthesized, voltage-sensitive fluorophore with a genetically encoded, self-labeling enzyme to enable voltage imaging in *Drosophila melanogaster*. Previously, we showed that a rhodamine voltage reporter (RhoVR) combined with the HaloTag self-labeling enzyme could be used to monitor membrane potential changes from mammalian neurons in culture and brain slice. Here, we apply this hybrid RhoVR-Halo approach *in vivo* to achieve selective neuron labeling in intact fly brains. We generate a *Drosophila* UAS-HaloTag reporter line in which the HaloTag enzyme is expressed on the surface of cells. We validate the voltage sensitivity of this new construct in cell culture before driving expression of HaloTag in specific brain neurons in flies. We show that selective labeling of synapses, cells, and brain regions can be achieved with RhoVR-Halo in either larval neuromuscular junction (NMJ) or in whole adult brains. Finally, we validate the voltage sensitivity of RhoVR-Halo in fly tissue via dual-electrode/imaging at the NMJ, show the efficacy of this approach for measuring synaptic excitatory post-synaptic potentials (EPSPs) in muscle cells, and perform voltage imaging of carbachol-evoked depolarization and osmolarity-evoked hyperpolarization in projection neurons and in interoceptive subesophageal zone neurons in fly brain explants following *in vivo* labeling. We envision the turn-on response to depolarizations, fast response kinetics, and two-photon compatibility of chemical indicators, coupled with the cellular and synaptic specificity of genetically-encoded enzymes, will make RhoVR-Halo a powerful complement to neurobiological imaging in *Drosophila*.

**Significance Statement:** Voltage imaging is a powerful method for interrogating neurobiology. Chemical indicators possess fast response kinetics, turn-on responses to membrane depolarization, and can be compatible with two-photon excitation. However, selective cell labeling in intact tissues and *in vivo* remains a challenge for completely synthetic fluorophores. Here, we show that a chemical – genetic hybrid approach in *Drosophila* enables cell-specific staining *in vivo* and voltage imaging in whole-brain explants and at neuromuscular junction synapses.

## Introduction

Voltage imaging in intact brains offers the tantalizing promise to watch, in real time, the electrical changes that underlie physiology. Approaches for voltage imaging rely on fluorescent indicators, either chemically synthesized, genetically encoded, or combinations of the two. Chemically-synthesized indicators have a storied past, but suffer from combinations of low sensitivity, slow response kinetics, and the inability to localize to defined neurons. More recently, genetically-encoded indicators of voltage changes circumvent problems of localization to specific neurons. However, genetically encoded indicators also face problems of localization at the cellular membrane, slow response kinetics, low brightness, turn-off or non-linear responses to voltage changes, and, for opsin-based indicators, incompatibility with two-photon (2P) illumination.

Our group has focused on the development of chemically-synthesized voltage-sensitive fluorophores that respond to changes in membrane potential via a photoinduced electron transfer (PeT) based mechanism. At hyperpolarizing potentials, the voltage across the membrane accelerates PeT from one side of the molecule to the other, short-circuiting and quenching fluorescence.^1^ At depolarized potentials, PeT is slowed, and the quantum yield of the dye increases. This configuration allows fast,^2^ linear, turn-on responses to depolarizations (with corresponding fluorescence decreases for hyperpolarization), good signal to noise, and compatibility with 2P excitation.^3-4^ However, attempts to deploy voltage-sensitive fluorophores in brain tissues resulted in comprehensive staining of all neuronal membranes, making it difficult to visualize clear boundaries between cells or regions of the brain.^3, 5^ Therefore, there is strong interest in developing hybrid systems in which voltage-sensitive dyes are directed to cells of interest, either via expression of exogenous enzymes^6-8^ or via targeting of native ligands.^9^ Other strategies involve targeting synthetic fluorophores to genetically-encoded voltage-sensitive proteins, whether opsins^10-11^ or voltage-sensing domains.^12^

We recently reported a chemical-genetic hybrid, in which a chemically-synthesized rhodamine-based voltage reporter (RhoVR)^13^ attached to a flexible polyethyleneglycol (PEG) linker terminating with a chloroalkane forms a covalent bond with a cell-expressed HaloTag (**Figure 1**), enabling voltage imaging from defined neurons, in mouse cortical brain slices.^14^ This approach, RhoVR-Halo, takes advantage of the fast kinetics, linear turn-on response, and 2P compatibility of RhoVR-type indicators,^3-4^ and pairs it with the ability to target specific cells using traditional genetic methods.

**Figure 1.**
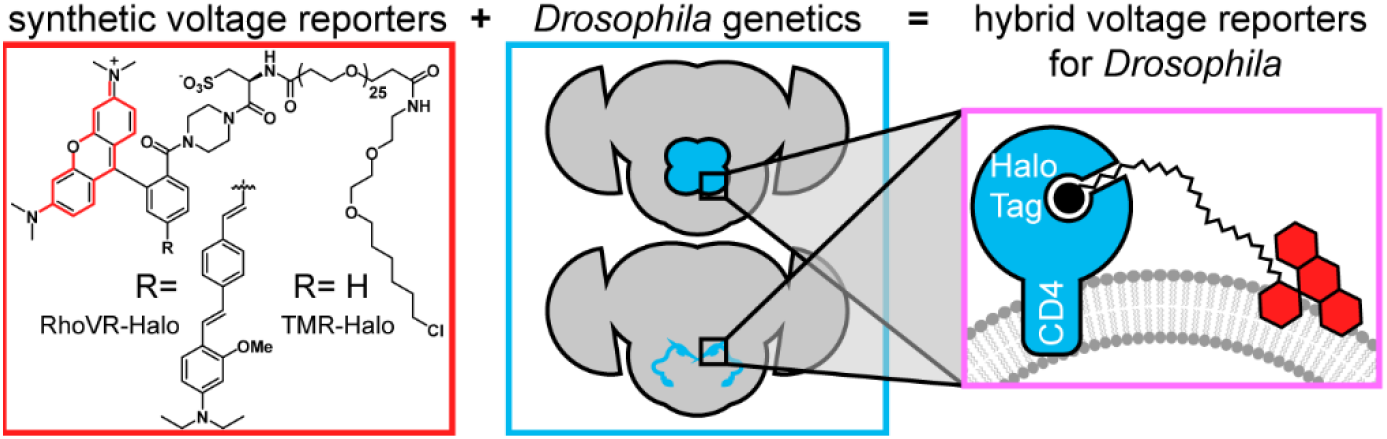
Chemical-genetic hybrids for voltage imaging in *Drosophila*. (red box) Chemically synthesized rhodamines with chloroalkane ligands will form covalent adducts with HaloTag enzymes. When R = the indicated molecular wire, the resulting RhoVR-Halo is voltage-sensitive. When R = H, the tetramethyl rhodamine-Halo is not voltage-sensitive (TMR-Halo). (teal box) The use of GAL4-UAS fly lines enables selective expression of HaloTag enzymes in defined populations of neurons. (magenta box) When HaloTag is fused with CD4, expression on the cell surface of defined neurons allows *in vivo* labeling (with either TMR-Halo or RhoVR-Halo) followed by *ex vivo* voltage imaging (with RhoVR-Halo).

The wealth of genetic tools, small brain size for optical imaging, and short generation time make *Drosophila melanogaster* an attractive model organism.^15-17^ Genetically encoded indicators have been previously deployed in *Drosophila* and fall into two broad classes: 1) fluorescent protein (FP) fusions with voltage-sensing domains and 2) electrochromic FRET indicators (eFRET) that couple voltage-dependent changes in opsin absorbance with FRET to a fluorescent protein. FP-VSD fusions like ArcLight^18-20^ or ASAP,^21-22^ have been used in multiple *Drosophila* contexts and show negative-going responses to membrane depolarizations, use “GFP”-like excitation and emission, and display non-linear response kinetics across the entire physiological range. Electrochromic-FRET indicators^23^ like Ace2N-mNeon^24^ or Varnam^25^ (and their chemigenetic relative, Voltron, which replaces the FP with a HaloTag)^10^ have also been used in *Drosophila* and provide fast, negative-going responses to depolarizations. These types of indicators are not compatible with 2P excitation, likely owing to the complex photocycle involved in opsin-based voltage sensitivity.^26^

Therefore, to expand the RhoVR-Halo methodology beyond vertebrate systems, we developed a stable transgenic UAS reporter line in *Drosophila* to express HaloTag on the extracellular surface of neurons, enabling the selective staining of defined neuronal populations when crossed with existing GAL4 driver lines. When paired with voltage-sensitive RhoVR-Halo,^14^ HaloTag-expressing flies allow cell type-specific labeling *in vivo*, and voltage imaging in a variety of contexts, including synaptic imaging at the neuromuscular junction (NMJ) and across multiple neurons in fly brain explants.

## Results

### Generation of HaloTag constructs for expression in flies

Although HaloTag and other self-labeling enzymes have been successfully expressed in transgenic flies, the reported lines localize HaloTag intracelluarly.^10, 27-28^ Our first task was to generate a HaloTag that expressed on the extracellular face of membranes. Previous chemical-genetic hybrids deployed in mammalian cells used a transmembrane domain from the platelet-derived growth factor receptor (PDGFR) to localize HaloTag to the cell surface and a secretion signal peptide from immunoglobulin K (IgK) to enhance export of the construct.^14^ To adapt HaloTag-directed chemical-genetic hybrids for voltage imaging in *Drosophila*, we selected CD4 as a transmembrane anchor, on account of its good membrane association in *Drosophila* neurons,^29^ fusing it to the C-terminus of the HaloTag. We sub-cloned this construct into different vectors for expression in mammalian (pcDNA3.1) and insect cells (pJFRC7).^30^

HaloTag-CD4 shows good expression on cell surfaces. In mammalian cells, CD4 localizes to the cell surface by anti-CD4 immunocytochemistry (**Figure S1**). Inclusion of the self-labeling enzyme, HaloTag, affords the opportunity to confirm not only localization, but activity of the expressed enzyme by delivering HaloTag substrates. HEK cells expressing HaloTag-CD4 and treated with RhoVR-Halo (100 nM) show good membrane localization (**Figure 2a**), while cells that do not express HaloTag-CD4 show approximately 3.5-fold lower fluorescence levels (**Figure 2b-c**). RhoVR-Halo survives fixation: following live-cell imaging, cells can be fixed and retain their RhoVR-Halo staining, which serves as a useful counterstain to the anti-CD4 immunocytochemistry (**Figure S1**). Live cells labeled with RhoVR-Halo and subsequently fixed, permeabilized with detergent and assayed for CD4 via immunochemistry reveal the majority of CD4 is found intracellularly, however RhoVR-Halo primarily localizes to cell membranes (**Figure 2b**). HEK cells expressing HaloTag-CD4 and labeled with TMR-Halo show approximately 15-fold greater fluorescence than cells that do not express HaloTag-CD4 (**Figure S2**).

**Figure 2.**
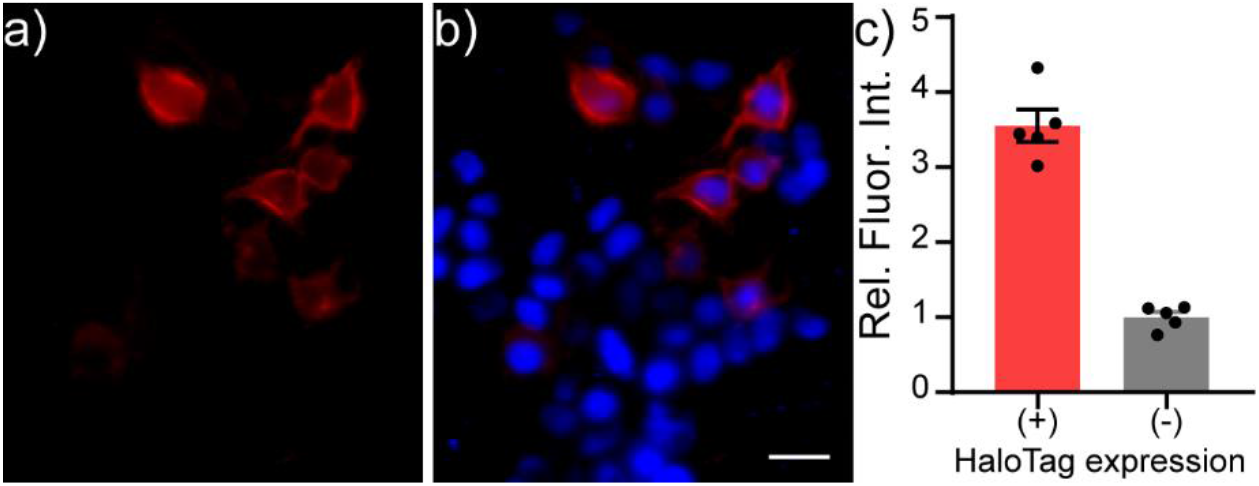
Live-cell staining of RhoVR-Halo in HEK293T cells expressing HaloTag-CD4. Epifluorescence images of HEK293T cells expressing HaloTag-CD4 (CMV promoter) and stained with **a)** RhoVR-Halo (100 nM, red) and **b)** Hoechst 33342 (1 μM, blue). Scale bar is 20 μm. **c)** Plot of relative fluorescence intensity in cells expressing HaloTag vs. cells that do not express HaloTag. HaloTag (+) cells were assigned based on a threshold obtained from a non-transfected control. Data are mean ± SEM for n = 5 different coverslips of cells. Data points represent average fluorescence intensities of 30 to 40 cells.

In S2 cells, an immortalized *Drosophila* cell line, we also observe cell surface localization of HaloTag-CD4, as visualized by anti-CD4 immunocytochemistry (**Figure S3**). S2 cells show similar HaloTag-CD4 dependent staining with TMR-Halo (100 nM, **Figure 3, Figure S3a-b**) with a 20-fold enhancement in fluorescence intensity in HaloTag-CD4 expressing cells compared to non-expressing cells (**Figure 3c**). TMR-Halo staining in S2 cells is also retained post-fixation (**Figure S3**).

**Figure 3.**
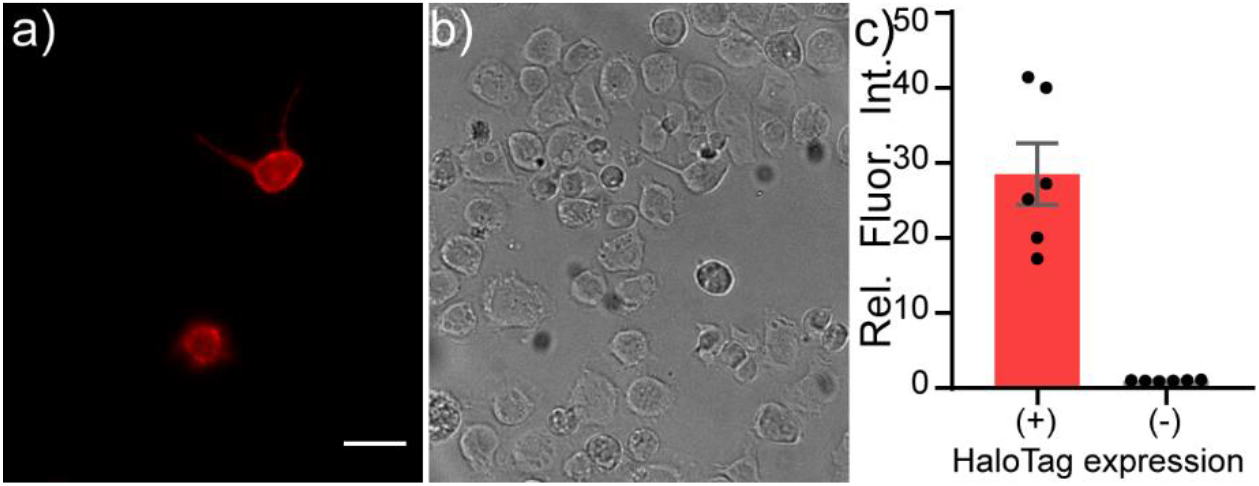
Live-cell staining of *Drosophila* S2 cells with TMR-Halo. Live-cell staining of TMR-Halo in *Drosophila* S2 cells expressing HaloTag-CD4. Epifluorescence images of *Drosophila* S2 cells transfected with tubP-GAL4 and HaloTag-CD4 UAS and **a)** stained with TMR-Halo (100 nM). **b)** Transmitted light image of cells in panel (**a**). Scale bar is 20 μm. **c)** Plot of relative fluorescence intensity in cells expressing HaloTag vs. cells that do not express HaloTag from the same cultures. HaloTag-(+) cells were assigned based on a threshold obtained from a non-transfected control. Data are mean ± SEM for n = 6 different coverslips.

We evaluated the voltage sensitivity of RhoVR-Halo in HaloTag-CD4 expressing HEK293T cells (**Figure 4**). After loading cells with RhoVR-Halo (500 nM), cells were subjected to whole-cell, patch-clamp electrophysiology. The voltage sensitivity of RhoVR-Halo in HaloTag-CD4 expressing HEK293T cells is approximately 14% per 100 mV (±2%, SEM n = 7 cells). This is approximately 70% of the value we obtained when HaloTag is targeted with previously developed^14^ HaloTag-pDisplay (**Figure S4**).

**Figure 4.**
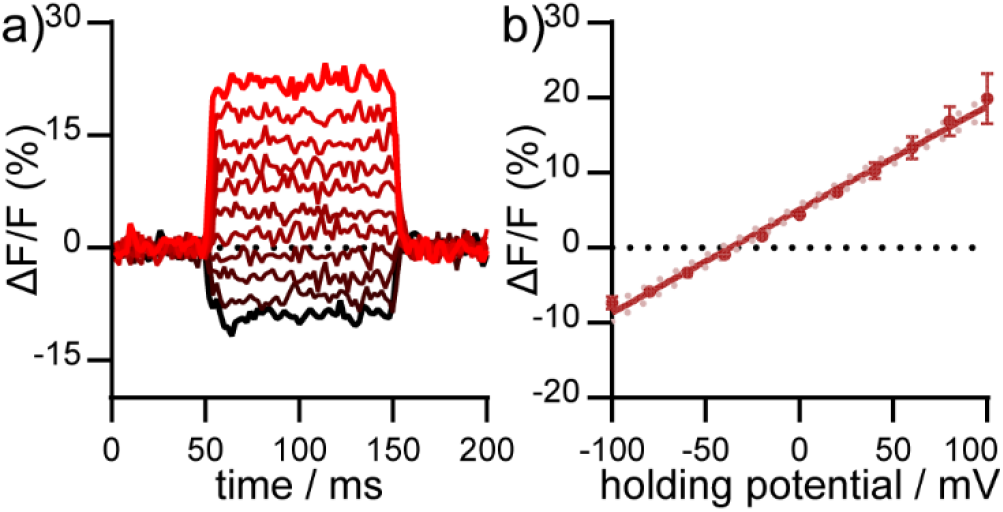
Voltage sensitivity of RhoVR-Halo in HEK293T cells expressing HaloTag-CD4. **a)** Plot of ΔF/F vs. time for a single HEK293T cell expressing HaloTag-CD4 and stained with RhoVR-Halo. The HEK293T cell was held at -60 mV and then stepped through hyperpolarizing and depolarizing potentials, in 20 mV increments, from -100 mV to +100 mV. **b)** Plot of ΔF/F vs. potential in mV. Data are mean ± standard error of the mean for n = 7 separate cells. Solid line is the line of best fit, and pink dots are 95% confidence interval.

### Validation of HaloTag-expressing Flies

To evaluate the performance of cell surface-expressed HaloTag-CD4 in intact flies, we generated transgenic flies (BestGene Inc.) and crossed the resulting UAS-HaloTag-CD4 line with a pan-neuronal driver line, neuronal synaptobrevin-GAL4 (nSyb-GAL4),^31^ which was used to drive HaloTag-CD4 expression in all neurons, (**Figure 5a**). Brains of nSyb-GAL4>HaloTag-CD4 flies show strong CD4 expression (**Figure 5a-c**). The pattern of anti-CD4 fluorescence indicates good localization to the plasma membrane (**Figure 5d-e**).

**Figure 5.**
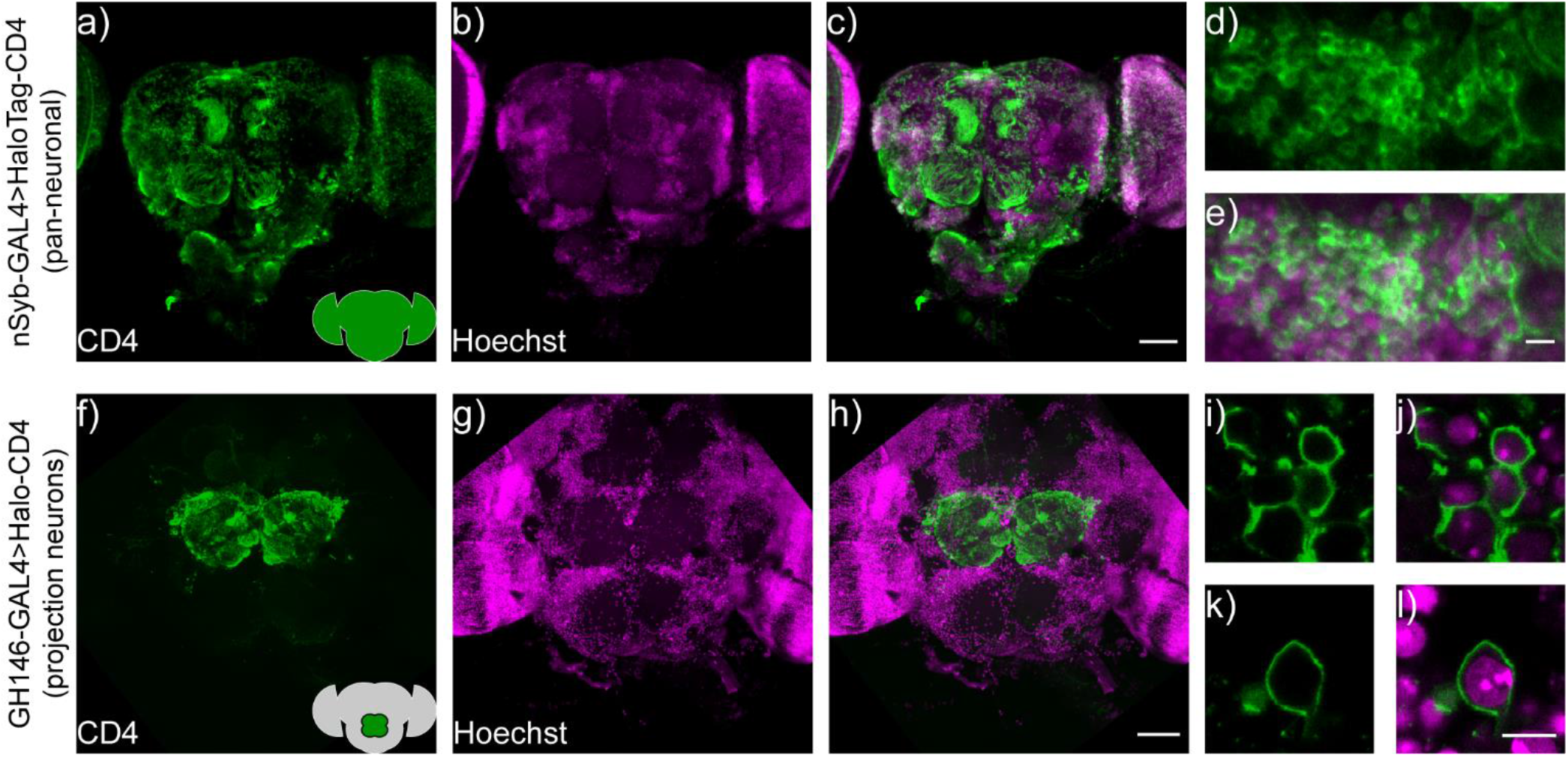
HaloTag-CD4 expression in transgenic *Drosophila*. **a)** nSyb-GAL4, HaloTag-CD4 brains express CD4 pan-neuronally. Maximum z-projection of a confocal fluorescence microscopy stack of brain explant from either **a-e)** nSyb-GAL4>HaloTag-CD4 or **f-l)** GH146-GAL4>HaloTag-CD4, fixed and stained for an extracellular epitope of the CD4 protein (OKT4, green) and counterstained for nuclei with Hoechst 33342 (16 μm or 10 μg/mL, magenta). Scale bar is 50 μm for whole-brain images (**a-c** and **f-h**) and 5 μm for zoomed-in regions (**d-e** and **i-l**). Insets on panels **a** and **f** show schematized brains with an approximate location of the staining for reference.

To evaluate the specificity of labeling, we expressed UAS-HaloTag-CD4 in a subset of neurons. We crossed UAS-HaloTag-CD4 flies with GH146-GAL4 flies^32-33^ to drive expression in a subpopulation of olfactory projection neurons (PNs) in the antennal lobe, a key olfactory relay. Immunohistochemistry reveals strong CD4 staining, localized to the antennal lobe in transgenic GH146-GAL4>HaloTag-CD4 flies (**Figure 5f-h**). These neurons also showed good extracellular staining (**Figure 5i-l**).

HaloTag remains functional when expressed on the cell surface of *Drosophila* neurons, enabling a range of brain regions and neurons to be labeled with small molecules. We delivered TMR-Halo (1 μM) to live flies via application of a solution of TMR-Halo in artificial hemolymph (AHL) to flies with their cuticle removed^34^ (see **Supplementary Info** for dissection details). We then imaged via confocal microscopy to establish the extent of labeling (**Figure 6**). In GH146-GAL4>HaloTag flies (PN labeling) treated with TMR-Halo, we observe strong fluorescence localized to the antennal lobe **(Figure 6a**). Non-transgenic fly controls show low fluorescence levels in the brain and antennal lobe (GH146-GAL alone, **Figure 6b**). TMR-Halo in combination with HaloTag-CD4 can be used to label single cells. VT011155-GAL4>HaloTag-CD4 fly brains drive expression in single interoceptive subesophageal zone neurons (ISNs),^19^ and treatment with TMR-Halo results in bright fluorescence localized to these neurons (**Figure 6c**). Similar staining profiles can be achieved with the voltage-sensitive RhoVR-Halo, which clearly labels PNs of the antennal lobe (**Figure 6d**, GH146-GAL4). High magnification examination of labeled projection neurons reveals membrane-localized staining (**Figure 6e** and **g**, red) alongside Hoechst 33342 nuclear staining (**Figure 6f** and **h, blue**). RhoVR-Halo can also label smaller sub-sets of neurons cells; treatment of Nan-GAL4>HaloTag-CD4 brains with RhoVR-Halo results in labeling of ISNs (**Figure 6i**).

**Figure 6.**
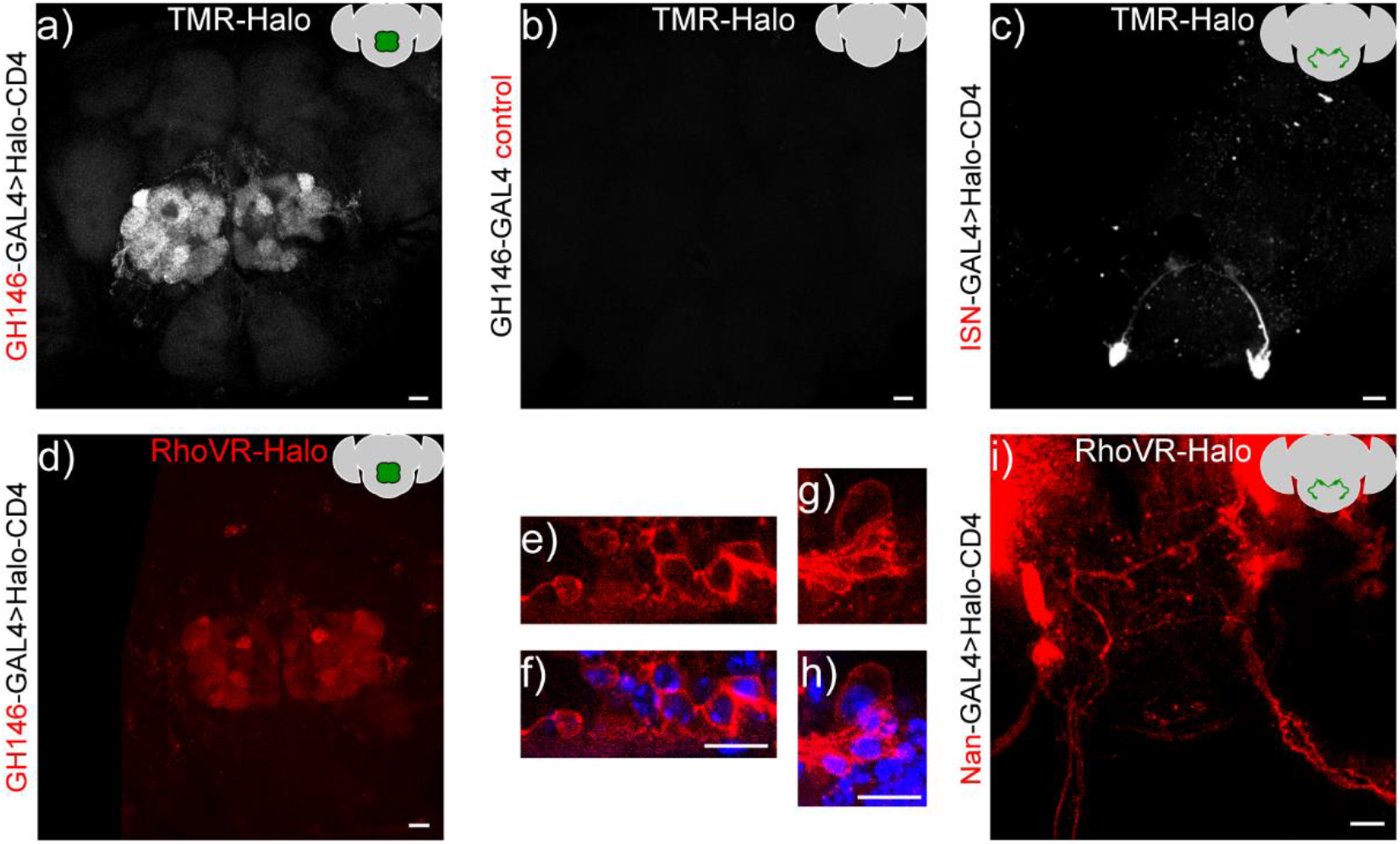
*In vivo* labeling of *Drosophila* neurons with TMR-Halo or RhoVR-Halo. Top row: Maximum z-projection of a confocal fluorescence microscopy stack of live brain explants labeled with voltage-*insensitive* TMR-Halo (1 μM) in an intact, live-fly before dissection and imaging. Crosses were either **a)** GH146-GAL4>HaloTag-CD4, **b)** GH146-only control, or **c)** VT011155-GAL4>HaloTag-CD4. Bottom row. Maximum or sum z-projections of confocal fluorescence microscopy stack of live brain explants labeled with voltage-*sensitive* RhoVR-Halo (1 to 2 μM), labeling either **d-h)** projection neurons (GH146-GAL4>Halo-CD4, max projection) or **i)** ISNs (Nan-GAL4>Halo-CD4, sum projection). **e** and **g)** High magnification images of RhoVR-Halo staining in PNs (red), overlaid with **f** and **h)** Hoechst 33342 nuclear stain (blue). All scale bars are 20 μm.

We used the same live-animal staining procedure to optimize the loading of RhoVR-Halo (**Figure S5**). We find that 2 μM RhoVR-Halo provides good staining in the antenna lobes of GH146-GAL4>HaloTag-CD4 crosses (**Figure S5**). Fluorescence from RhoVR-Halo is localized to the periphery of cell bodies, again supporting the extracellular expression of HaloTag-CD4 (**Figure 6e-h**). Compared to regions of the brain that do not express HaloTag-CD4, RhoVR-Halo fluorescence is approximately 3 times higher (**Figure S5b-c**). We find homozygous flies for GH146-GAL4>HaloTag-CD4 have slightly higher fluorescence levels compared to levels of heterozygous flies, when stained with the same concentration of RhoVR-Halo (**Figure S5h-j**). However, because the difference in fluorescence intensity in homozygous flies was not significantly larger than heterozygotes, we used heterozygous flies for subsequent experiments.

### Functional Imaging

We established the voltage sensitivity of RhoVR-Halo in fly tissue expressing HaloTag-CD4 using two different approaches. First, we performed electrophysiology using dual two-electrode voltage-clamp combined with fluorescence imaging at the larval *Drosophila* neuromuscular junction (NMJ). We used the motor neuron driver OK6-GAL4 to drive pre-synaptic expression of HaloTag-CD4 (**Figure 7a-f**) or the muscle driver G14-GAL4 to express HaloTag-CD4 in the post-synaptic muscle (**Figure 7g-l**). In live 3^rd^ instar larval NMJ preparations, RhoVR-Halo (2 μM) clearly stains pre-synaptic neuronal compartments when HaloTag-CD4 expression is targeted in motor neurons (red, **Figure 7d**), co-localizing with the neuronal plasma membrane marker horseradish peroxidase (HRP, grey, **Figure 7e-f**). In a complementary fashion, when HaloTag-CD4 is expressed in post-synaptic muscle cells, RhoVR-Halo fluorescence (red, **Figure 7j**) accumulates at NMJs outside of the neuronal membrane outlined by HRP (grey, **Figure 7k-l**). RhoVR-Halo readily detects excitatory post-synaptic potentials (EPSPs) in muscle cells, confirmed by simultaneous optical imaging and sharp electrode recordings (**Figure 7m-o**). Importantly, we next used two-electrode voltage-clamp recordings in a semi-dissected larval preparation with muscle HaloTag-CD4 expression (G14-GAL4>HaloTag-CD4, **Figure 7p**). This approach demonstrated that depolarizing potentials result in an increase in RhoVR-Halo signal (m6, **Figure 7q-s**) with an overall voltage sensitivity of approximately 12% ΔF/F per 100 mV (± 0.2%, n = 8), in reasonably close agreement to the value determined in HEK293T cells (14%, **Figure 4**). Analysis of electrode (**Figure 7r**) and optical recordings (**Figure 7s**) show good correspondence. In contrast, no change in fluorescence signals was observed in an adjacent unclamped/unstimulated muscle cell (m7, **Figure 7q**, grey).

**Figure 7.**
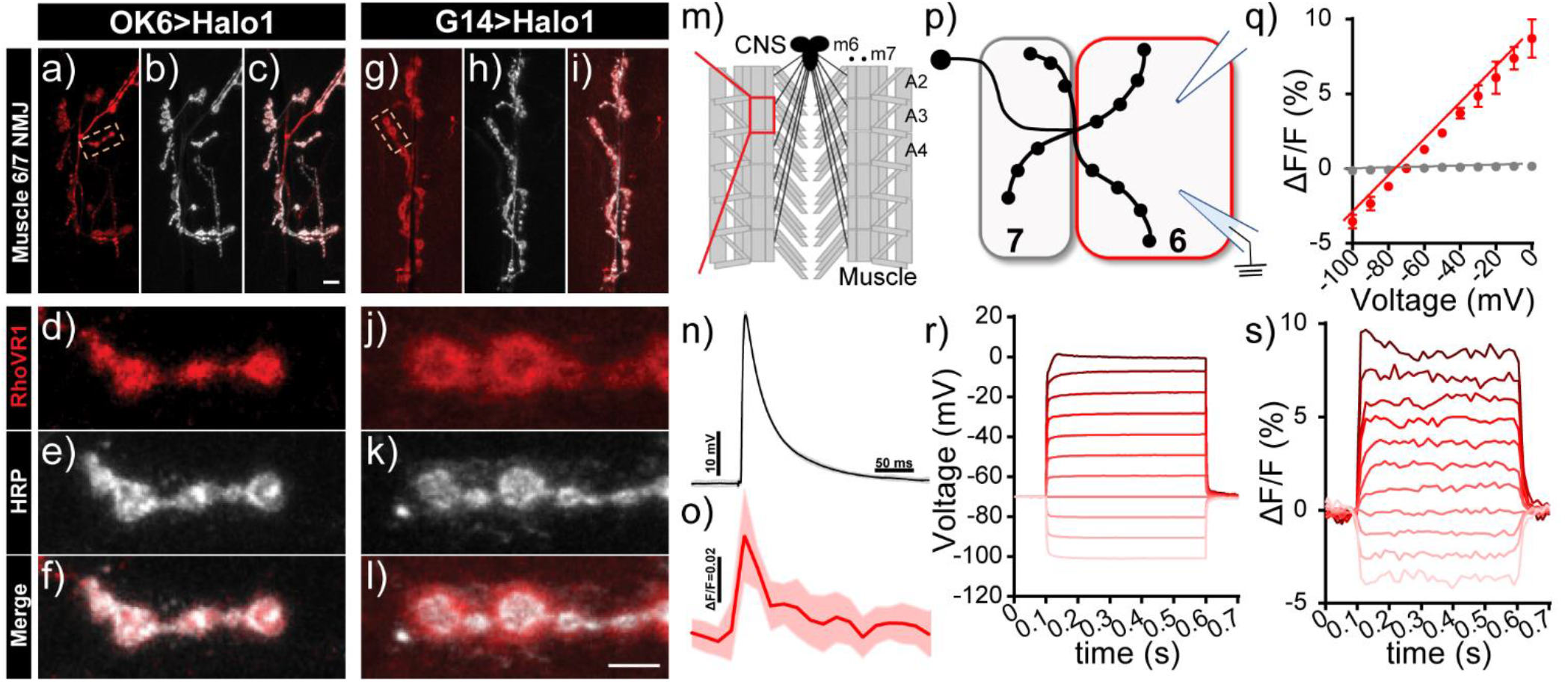
Voltage imaging with RhoVR-Halo using the *Drosophilia* neuromuscular junction (NMJ). **a-l)** Confocal images of motor neurons labeled with RhoVR-Halo (2 μM) in NMJs of **a-f)** presynaptic neuron-labeled OK6-GAL4>Halo-CD4 flies or **g-l)** post-synaptic muscle-labeled G14-GAL4>Halo-CD4 flies. Red is RhoVR-Halo fluorescence; grey is HRP – a neuronal membrane marker. Scale bars are 10 μm (a-c, g-i) and 5 μm (d-f, j-l). **m)** Schematic of *Drosophila* NMJ. Excitatory post-synaptic potentials (EPSPs) recorded at NMJs of G14-GAL4>Halo-CD4 larvae stained with RhoVR-Halo (2 μM). Sharp electrode recordings of EPSPs are in **n)** grey, and **o)** optically recorded EPSPs are in red. Data are mean ± SEM of 8 replicates. **p)** Schematic of two-electrode measurements. Muscle cell 7 is unclamped, while the membrane potential of muscle cell 6 (m6) is clamped, held at -70 mV, and stepped to hyper- and depolarizing potentials ranging from -100 mV to 0 mV. **q)** Plot of ΔF/F vs. holding potential for m6 (clamped, red) or m7 (unclamped, grey) in G14-GAL4>Halo-CD4 flies stained with RhoVR-Halo. Data are mean ± standard error of the mean for n = 8 independent determinations. Example plots of change in **r)** voltage or **s)** fluorescence (ΔF/F) vs. time for the clamped m6 cell.

As a second confirmation of voltage sensitivity in fly tissues, we developed a stereotyped stimulation protocol for imaging in fly brain explants. We generated flies that express both HaloTag and the voltage-sensitive fluorescent protein, ArcLight, in PNs (GH146 GAL4, HaloTag/CyO; ArcLight/TM2) for use as an internal positive control. The use of RhoVR-Halo, with excitation and emission profiles in the green/orange region of the visible spectrum, allows for the simultaneous deployment of GFP-based indicators,^13-14^ like ArcLight.^18^ *Drosophila* antennal lobe projection neurons receive input from the olfactory receptor neurons (ORNs) in the antennae.^35^ As these projection neurons primarily receive cholinergic input from the ORNs,^36^ we hypothesized that PNs could be readily stimulated with carbachol (CCH), a non-hydrolyzable acetylcholine mimic. We treated ArcLight/HaloTag-CD4 expressing fly brain explants with carbachol (100 μM) and observed robust fluorescence decreases timed to carbachol treatment, indicating a depolarizing membrane potential response to this neurotransmitter analog (**Figure S6a-d**).

Using this robust stimulation protocol in fly brain explants, we next performed two-color voltage imaging with RhoVR-Halo and ArcLight. As before, we loaded RhoVR-Halo (2 μM) in live flies, removed the brains, and imaged the brain explants using epifluorescence microscopy. Excitation provided alternately with blue (475 nm) or green (542 nm) light to excite ArcLight or RhoVR-Halo, respectively, revealed robust fluorescence responses to carbachol (100 μM) treatment (**Figure 8**). RhoVR-Halo fluorescence increases with carbachol stimulation (**Figure 8a-d**), corresponding to membrane voltage depolarization and the turn-on response of RhoVR-type indicators.^13-14^ In contrast, ArcLight fluorescence decreases with carbachol stimulation (**Figure 8e**), showing a fluorescence decrease in response to depolarization, consistent with the turn-off response to depolarization for ArcLight indicators.^18^ Importantly, neither RhoVR-Halo nor ArcLight responds to a control experiment that omits carbachol from the perfusion solution (**Figure 8c and f**). Finally, the chemical-genetic hybrid approach of RhoVR-Halo enables additional controls to be carried out using the same transgenic flies. When HaloTag/ArcLight expressing flies are treated with TMR-Halo and then stimulated with carbachol, there is no response from the voltage-insensitive TMR-Halo (**Figure 8c and g**), but ArcLight still responds (**Figure 8g**). Using a “functionally dead” rhodamine dye in this experiment allows for control experiments to be run in the same transgenic animals as the experiments. Similar experiments with inactive mutants of genetically-encoded indicators/actuators (like ArcLight or GCaMP) would require the generation of separate transgenic animals with the inactivating mutation.

**Figure 8.**
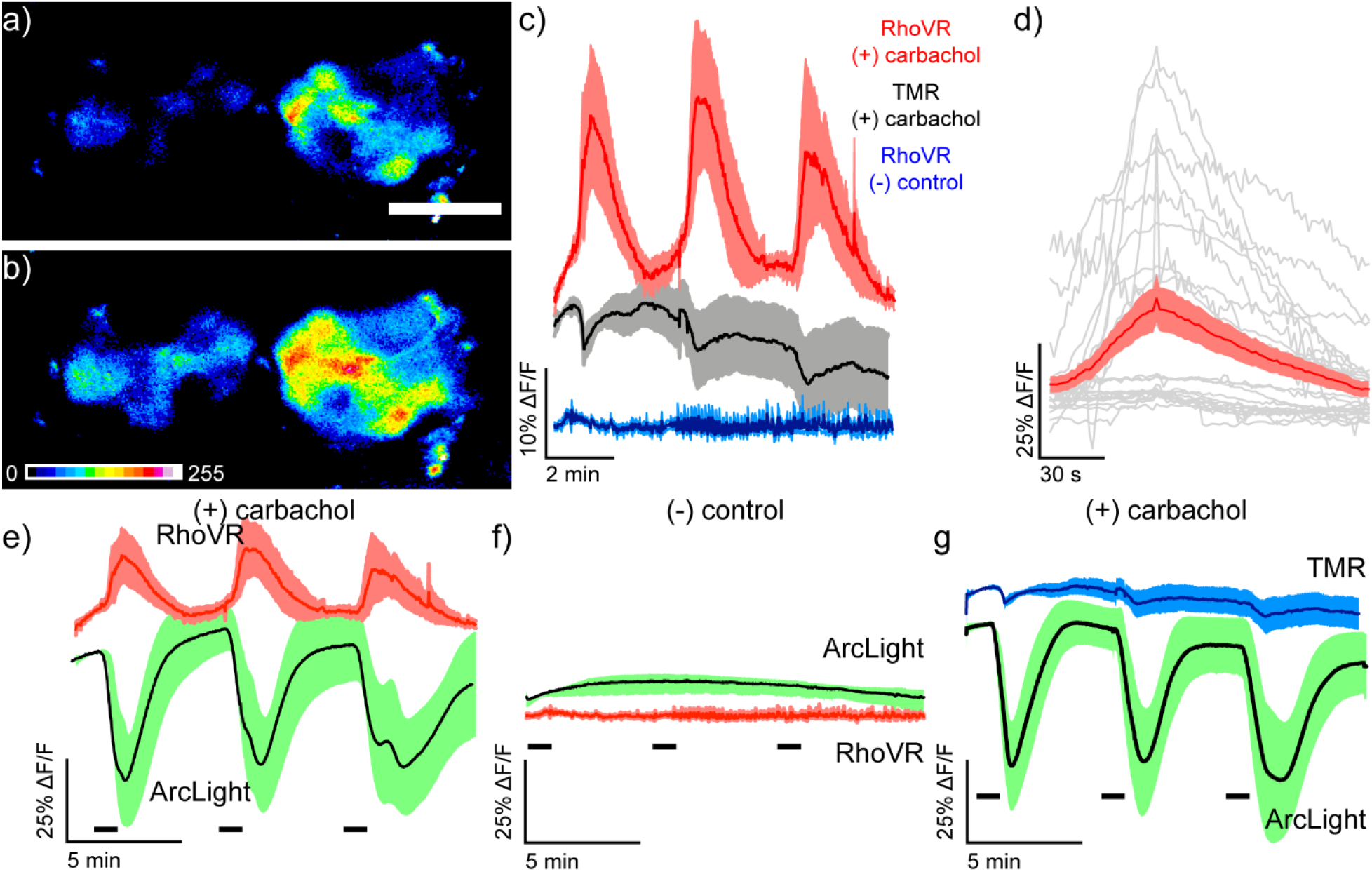
Simultaneous two-color visualization of carbachol-induced depolarization in projection neurons of live *Drosophila* brain explants with RhoVR-Halo and ArcLight. Epifluorescence images of live explant *Drosophila* brain expressing HaloTag-CD4 in antennal lobe projection neurons (GH146-GAL4, UAS-HaloTag-CD4/CyO; UAS-ArcLight/TM2) and labeled with RhoVR-Halo (2 μM) in live flies before dissection and explant imaging **a)** immediately before and **b)** 30 s after stimulation with 100 µM carbachol. Scale bar is 50 μm. **c)** Plots of average ΔF/F traces for *Drosophila* brains under the following conditions: stained with voltage-sensitive RhoVR-Halo (2 μM) and stimulated with 100 μM carbachol (red, n = 7 brains), stained with voltage-insensitive TMR-Halo (100 nM) and stimulated with 100 μM carbachol (grey, n = 7 brains), or stained with voltage-sensitive RhoVR-Halo (2 μM) and treated with vehicle control (blue, n = 6 brains). **d)** Plots of individual ΔF/F responses from RhoVR-Halo to carbachol stimuli (grey) and the average across all responses (red, SEM in light red). Traces of responses were aligned by peak response time and display 50 seconds before peak response and 150 seconds after peak response (grey). **e)** Plots of average ΔF/F traces for *Drosophila* brains stained with voltage-sensitive RhoVR-Halo (2 μM) and stimulated with 100 μM carbachol (red, n = 7 brains). ArcLight responses are recorded simultaneously (green, n = 7 brains). RhoVR traces are replicated from panel (**c**) for comparison with ArcLight. **f)** Plots of average ΔF/F traces for *Drosophila* brains stained with voltage-sensitive RhoVR-Halo (2 μM) and then treated with a vehicle control (red, n = 6 brains). ArcLight responses are recorded simultaneously (green, n = 6 brains). RhoVR traces are replicated from panel (**c**) for comparison with ArcLight. **g)** Plots of average ΔF/F traces for *Drosophila* brains stained with voltage-*insensitive* TMR-Halo (100 nM) and stimulated with 100 μM carbachol (blue, n = 6 brains). ArcLight responses are recorded simultaneously (green, n = 6 brains). TMR traces are replicated from panel (**c**) for comparison with ArcLight. For all plots, data are mean ±SEM for the indicated number of samples. *Drosophila* brain explants were stimulated three times for 30 s with either 100 μM carbachol or vehicle. Stimulus (delivery of carbachol or vehicle) is depicted by small black bars immediately below the traces).

To evaluate the ability of RhoVR-Halo to report on physiological stimuli, we probed the response of RhoVR-Halo in ISNs, cells that respond dynamically to changes in osmolarity. Previous studies demonstrated that increases in osmolarity (240 mOsm to 440 mOsm) evoke hyperpolarizing responses in ISNs.^19^ Consistent with this, we find that ISNs expressing HaloTag-CD4 (Nanchung-GAL4) and labeled with RhoVR-Halo hyperpolarize upon an increase in osmolarity, as indicated by decreases in RhoVR fluorescence (**Figure 9a-c**). In fly brains labeled with voltage-insensitive TMR-Halo, we observe no change in fluorescence (**Figure 9a-c**). In contrast, flies expressing ArcLight in ISNs show fluorescence increases in response to increased osmolarity (**Figure S7**). Two-color voltage imaging alongside ArcLight in flies that express both HaloTag-CD4 and ArcLight in ISNs (Nanchung-GAL4, UAS-HaloTag-CD4/CyO; UAS-ArcLight/TM2) reveals osmolarity-induced decreases in RhoVR fluorescence coupled with increases in ArcLight fluorescence (**Figure 9d-f**), while control experiments at constant osmolarity show no responses in either ArcLight or RhoVR fluorescence (**Figure 9g**). Heterozygous flies expressing HaloTag in ISNs and labeled with RhoVR-Halo also respond to changes in osmolarity (**Figure S8**). Taken together, these data establish the utility of RhoVR-Halo for monitoring sensory-induced changes to membrane potential.

**Figure 9.**
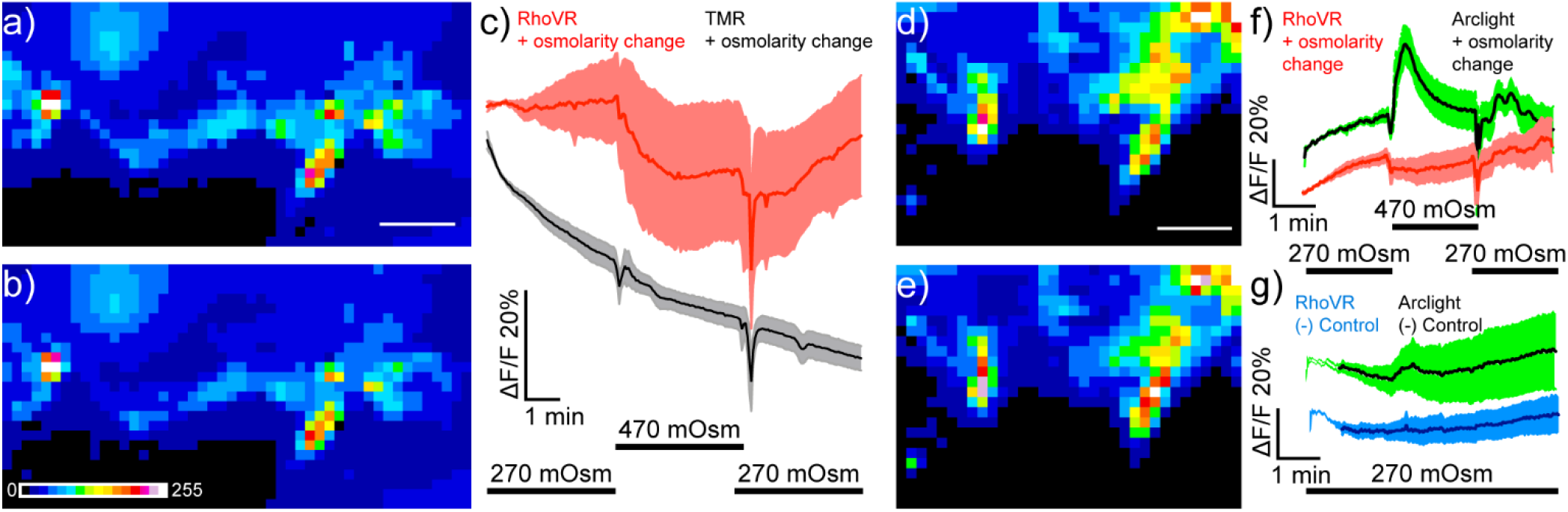
Imaging osmolarity induced hyperpolarizations in *Drosophila* interoceptive neurons in live explants using single color and dual-color imaging. Spinning disk confocal maximum z-projections of live explant *Drosophila* brain expressing HaloTag in ISNs (Nanchung-Gal4, UAS-HaloTag/Nanchung-Gal4, UAS-HaloTag; TM2/TM6B) **a)** before and **b)** after stimulation with high osmolarity hemolymph (470 mOsm). Scale bar is 50 μm. Image is pseudo-colored, and the scale bar indicates 8-bit pixel grey values. **c)** Plot mean of fluorescence (%ΔF/F) vs. time in response to one osmolarity simulation of either RhoVR-Halo (red, n = 5) or HT-TMR (black, n=5) loaded brains. Data are mean ± SEM Black bars below indicate the stimulation time course switching from 270 mOsm to 470 mOsm. Maximum z-projections of live explant *Drosophila* brain expressing HaloTag and Arclight in the ISNs (Nanchung-Gal4, UAS-HaloTag/UAS-Arclight; TM2/TM6B) **d)** before and **e)** after stimulation with high osmolarity hemolymph (470 mOsm). **f)** Plot mean of fluorescence (%ΔF/F) vs. time for simultaneously imaged Arclight (green) and RhoVR-Halo (red) in response to high osmolarity simulation (n=7) **g)** Plot mean of fluorescence (%ΔF/F) vs. time for simultaneously imaged Arclight (green) and RhoVR-Halo (red) in response a vehicle control (n=7).

## Discussion

In summary, we show that RhoVR-Halo indicators can be used for direct visualization of membrane potential changes in synapses and brains of flies. We show, for the first time, that RhoVR-Halo dyes can label specific neurons *in vivo* and that voltage changes can be visualized using epifluorescence microscopy at synapses in the NMJ and whole-brain explants. The hybrid chemical-genetic strategy employed here features a turn-on response to membrane depolarization and affords the opportunity to “plug-and-play” different fluorescent dyes to enable imaging in different colors^37^ or to run critical control experiments using a non-voltage-sensitive fluorophore in the same genetic background (**Figure 8g-i**). We envision that RhoVR-Halos, with their high two-photon (2P) cross-section (93 GM at 840 nm, **Figure S9**), can be combined with high-speed 2P imaging methods to provide fast voltage imaging in the brain.

Despite these advances, several drawbacks are associated with this methodology at present. First, in the imaging data presented here, we do not take full advantage of the response kinetics of PeT-based indicators like RhoVR, which should have nanosecond responses times based on the mechanism of voltage sensing.^2, 38-39^ Secondly, we do not take full advantage of the high 2P excitation cross-section of RhoVR dyes. Especially notable is the substantial cross-section at ∼1030-1040^4^ nm (**Figure S9**), which allows for the use of high-powered 2P illumination in emerging fast 2P methods.^4, 40^ Third, in fly brains, RhoVR-Halo voltage-sensitive indicators are not as bright as their fluorophore-only counterparts. This is likely a result of combinations of a) lower intrinsic quantum yield for RhoVR-Halo compared to TMR-Halo (since the presence of a molecular wire quenches the fluorescence of the dye) and b) lower solubility for the rather greasy RhoVR-Halo indicators compared to the smaller, more compact TMR-Halo dyes. The former can be addressed by using published methods to generate brighter fluorophores. The latter can be addressed by the use of new chemistries to attach HaloTag ligands, freeing up other sites for solubilizing groups. Even with these limitations, we envision that chemical-genetic hybrids like RhoVR-Halo will be an important complement to the expanding set of methods for visualizing membrane potential changes in living systems, especially in contexts where turn-on response to depolarization, fast kinetics, and 2P compatibility are required.

## Acknowledgments

We acknowledge support from NIH (R01NS098088, EWM; R01NS111414, DD) and NSF (NeuroNex Innovation Award 1707350, EWM and KS). MJK was supported, in part, by a training grant from the NIH (T32GM007232). BRB was supported, in part, by a training grant from the NIH (T32GM066698). MD acknowledges support from the NIH BRAIN Initiative (U24NS109107).

