## Supporting Information Document for "Voltage imaging in *Drosophila* using a hybrid chemical-genetic rhodamine voltage reporter"

**DOI:**

**Table of Contents**

General Methods ... 2
Plasmid construction. ... 2
Cell culture and transfection. ... 3
Dye loading. ... 4
Epifluorescence microscopy. ... 4
Epifluorescent image analysis. ... 5
Immunocytochemistry. ... 5
Transgenic generation. ... 6
Immunohistochemistry. ... 6
Live-fly brain dye loading with HT-TMR and voltage-sensitive dyes. ... 6
Confocal Microscopy. ... 7
Confocal imaging analysis. ... 8
Carbachol ArcLight functional imaging. ... 8
Dual Color Functional Imaging. ... 9
Functional imaging data analysis. ... 9
NMJ Imaging and Electrophysiology. ... 10
Two-photon excitation spectra and cross-sections. ... 11
Supporting Figures ... 14
Figure S1. Immunocytochemistry of HaloTag-CD4 in HEK293T cells. ... 14
Figure S2. TMR-Halo staining in HEK293T cells. ... 15
Figure S3. Immunocytochemistry in *Drosophila* S2 cells. ... 16
Figure S4. HaloTag. Comparison of voltage sensitivity of RhoVR-Halo with CD4-HaloTag and pDisplay-17
Figure S5. Optimization of dye loading in live-fly brains using voltage-sensitive dye RhoVR-Halo. 18

### General Methods

#### Plasmid construction.

We included a secretion signal derived from the signal peptide of the *Caenorhabditis elegans* β-integrin PAT-3[1] at the N-terminus of HaloTag, with the 5' UTR from *heat shock protein 70* (*hsp70*) and the 3'UTR and polyA tail from SV40 early genes, as described previously.[2]

For expression in HEK cells, we subcloned HaloTag via restriction digest (NheI, SalI) and subsequent Gibson Assembly into pCDNA3.1 vector containing a cytomegalovirus (CMV) promoter, a 5' PAT3 secretion signal, and a 3' CD4 transmembrane domain. For expression in S2 cells and transgenic generation, the insert Pat3-HaloTag-CD4 was assembled into pJFRC7[3] backbone via restriction digest (CD8::GFP was removed by XhoI and XbaI) and Gibson assembly (Addgene). All constructs were sequence confirmed by the UCB Sequencing Facility. Sequences used for all constructs can be found in the attached electronic construct maps.

**Scheme S1. Plasmid maps**

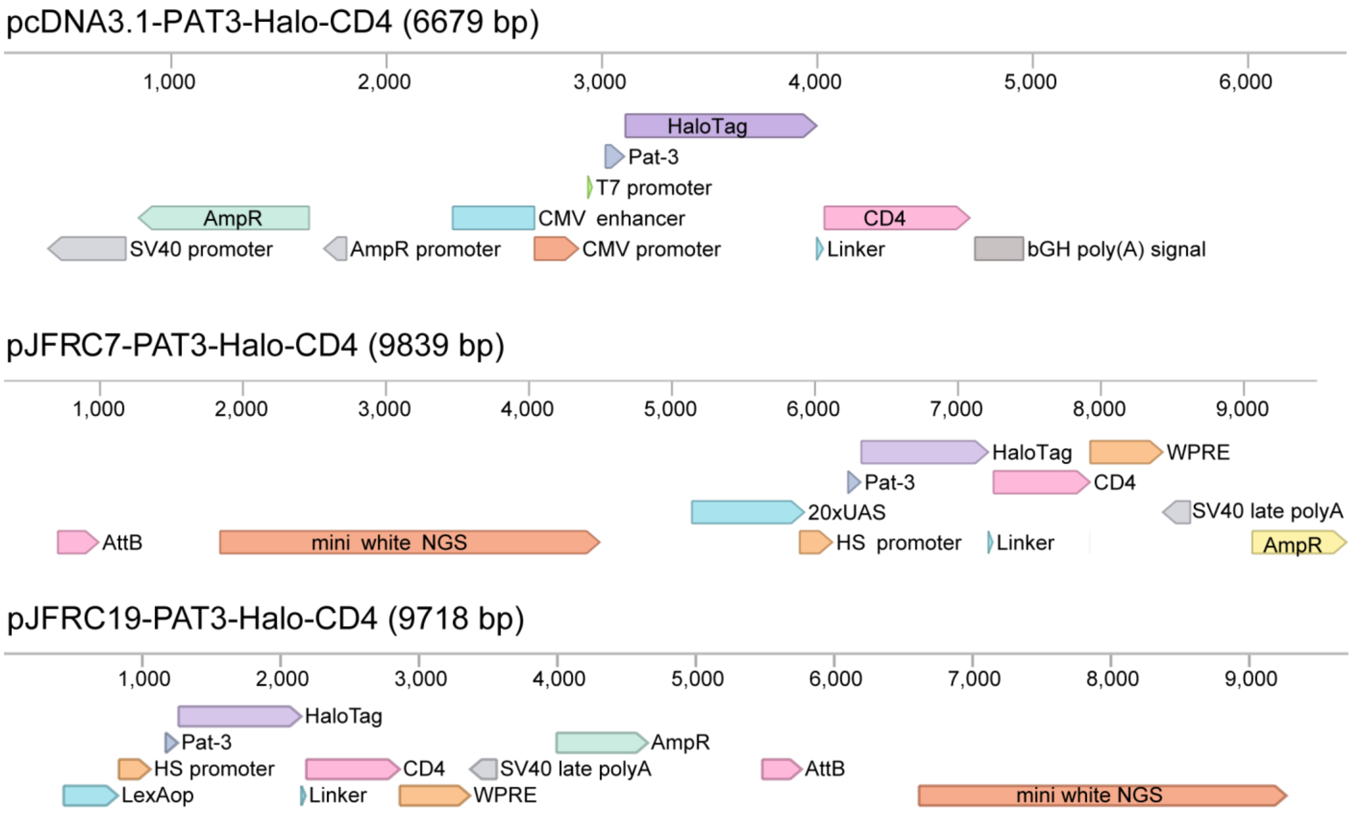

### Cell culture and transfection.

We obtained all cell lines from the UCB Cell Culture Facility. Human embryonic kidney 293T (HEK) cells were maintained in Dulbecco's modified eagle medium (DMEM) supplemented with 1 g/L D-glucose, 10% fetal bovine serum (FBS; Thermo Scientific), and 1% GlutaMax (Invitrogen) at 37 °C in a humidified incubator with 5 % CO2. Cells were passaged and plated in DMEM (as above) at a density of 50,000 cells onto 12 mm coverslips pre-treated with Poly-D-lysine (PDL;1mg/ml; Sigma-Aldrich). Transfection of plasmids was carried out using Lipofectamine 3000 (Invitrogen) 12 hours after plating. Imaging was performed 36 hours after plating.

S2 Cells were maintained in Schneider's Drosophila media (Thermo Fisher Scientific) supplemented with 10% FBS at 28°C in a non-humidified incubator under atmospheric conditions. Cells were passaged and plated at 500,000 cells per well in a 24 well plate. Six hours after plating, promotor Tubulin Gal4 pCaSper

(Addgene #17747)[4] was cotransfected with pJFRC7 constructs using a modified Lipofectamine 3000 (Life Technologies) protocol. This protocol included a 20-minute preincubation of lipofectamine and DNA in Opti-MEM (Life Technologies) and no p3000 reagent. Forty-eight hours after transfection, S2 cells were transferred onto PDL-treated (1 mg/mL) 12 mm coverslips and allowed to adhere for 30 minutes before dye loading and imaging.

**Dye loading.**

We maintained DMSO stock solutions (100 µM) of all dyes at -80 °C in single-use aliquots. Aliquots were further diluted to a working concentration of 100 nM in HBSS and incubated on cells for 30 minutes at 37 °C for HEK cells and room temperature for S2 cells. We then replaced all dye-containing HBSS with fresh HBSS and imaged in HBSS at room temperature.

**Epifluorescence microscopy.**

Imaging was performed on an AxioExaminer Z-1 (Zeiss) equipped with a Spectra-X Light engine LED light (Lumencor), controlled with Slidebook (v6, Intelligent Imaging Innovations). Images were acquired with a W-Plan-Apo 20x/1.0 water objective (20x; Zeiss) and focused onto an OrcaFlash4.0 sCMOS camera (sCMOS; Hamamatsu). The optical setup for imaging with each dye is described below.

**Table S1.** Optical filter sets for epifluorescence microscopy

| Dye | Excitation | Emission | Dichroic |
|---|---|---|---|
| HT-RhoVR1 | 525-560nm | 593/40 BP | 562 LP |
| HT-TMR | 525-560nm | 593/40 BP | 562 LP |
| A488 | 455-495nm | 430/32,<br>508/14, | 432/38,<br>509/22, |

| | | | |
|---|---|---|---|
| | | 586/30,<br>708/98 BP | 586/40, 654 nm LP |
| Hoechst 33342 | 375-400nm | 405/40 BP | 415 LP |

**Epifluorescent image analysis.**

For fluorescence intensity measurements, regions of interest were drawn around cell bodies, and fluorescence was calculated in ImageJ (FIJI), NIH). We identified transfected cells by setting a threshold that excluded all cells in the non-transfected controls. We calculated the fold change between non-transfected and transfected cells by taking the ratio of transfected cells fluorescence and non-transfected cell fluorescence. For each condition, at least 100 cells were circled across three to five individual coverslips.

**Immunocytochemistry.**

Immediately following live-cell dye loading experiments, cells were fixed for 20 minutes at room temperature with 4% formaldehyde in PBS. Cells were then washed in PBS (3x 5-minute washes) and treated with either 0.3% Triton X-100 in PBS for the permeabilized condition or PBS for the nonpermeabilized condition. Cells were again washed in PBS and blocked for at least 45 minutes in 0.1% NGS in PBS. Cells were then incubated overnight at 4 °C with 1:500 mouse anti CD4 (OKT4; Thermo Fisher Scientific). We then washed each sample in PBS and stained with a spectrally compatible mouse secondary Goat anti-Mouse A488 (Life Technologies) or Goat anti-Mouse A647 (Life Technologies) 1:1000 in 0.1%NGS for 2hrs at room temperature. We added Hoechst 33342 (10mg/mL -20 stock) 1:1000 for the last 15 minutes of this incubation period. Cells were then washed (3 x 5-minute washes) in PBS and mounted onto glass slides using Fluoromount Mounting Media (VWR International) before imaging.

**Transgenic generation.**

pJFRC7-Pat3-HaloTag-CD4 were sent to Best Gene Inc. for injection into the following genomic sites via phi-C31 integration.

**Table S2.** Injection phi C31 site and stock line

| Construct | Injection Site | Injection Stock |
|---|---|---|
| pJFRC7-Pat3-HaloTag-CD4 | VIE260B | VDRC#60100 |

**Immunohistochemistry.**

Fly brains were dissected in calcium-magnesium free artificial hemolymph (AHL-/-; NaCl 108.0 mM, KCl 5.0 mM, $NaHCO_3$ 4.0 mM, $NaH_2PO_4 \cdot H_2O$ 1.0 mM, Trehalose· 2 $H_2O$ 5.0 mM, Sucrose 10.0 mM, HEPES 5.0 mM and adjusted to pH 7.5 with NaOH) and fixed for 20 minutes in 4% formaldehyde in PBS. Brains were then washed in PBS (3x 5-minute washes) and treated with either 0.3% Triton x -100 in PBS for the permeabilized condition or PBS for the nonpermeabilized condition. Brains were again washed in PBS and blocked for at least 45 minutes in 0.1% NGS in PBS. Cells were incubated overnight in block containing 1:50 RT anti HA (Sigma Aldrich) or 1:100 or 1:200 mouse anti CD4 at 4 °C for 48 hours. Brains were then washed in PBS (3 x 5-minute washes) and stained with Goat anti-Rat A488 (Life Technologies) 1:1000 and Goat anti-Mouse A594 (Life Technologies) in block for 4 hours at room temperature shaking. We added Hoechst 33342 (10 mg/mL) 1:1000 for the last 15 minutes of this incubation. Brains were then washed and mounted onto glass slides using Vectashield mounting media (Vector Laboratories) before imaging using confocal microscopy.

**Live-fly brain dye loading with HT-TMR and voltage-sensitive dyes.**

Live-fly preparations were performed in the following way: 10-40 day old flies were briefly anesthetized on ice and placed into a small slit on a custom-built plastic mount at the cervix so that the head was isolated from the rest of the body. The head was then immobilized using clear nail polish, which was allowed to set for 15 minutes. The head cuticle was then removed using sharp forceps in calcium-magnesium free Artificial Hemolymph solution (AHL-/-)[5], and the esophagus was cut to eliminate autofluorescence. The AHL was then replaced with calcium-magnesium free AHL containing 0.2% Pluronic F127, and either 2 µM RhoVR-Halo or 1 µM TMR-Halo (for functional imaging experiments, this was lowered to 100 nM TMR-Halo to match fluorescence intensity with RhoVR-Halo), and the glial sheath was punctured manually over the optic lobes to permit dye access. DMSO concentrations were maintained below 3% vol/vol in the dye loading solutions. Following a 15-minute loading period in the dark at room temperature, the brains were removed and then imaged via confocal or epifluorescent microscopy. When imaged under confocal microscopy, brains were mounted onto glass coverslips with spacers to prevent sample loss and deformation. When imaged under epifluorescent microscopy, brains were adhered to PDL- (Poly-D-Lysine), or PLL- treated (Poly-L-Lysine) coverslips and bathed in AHL.

**Confocal Microscopy.**

We performed confocal imaging using an LSM710 upright confocal microscope maintained by the Biological Imaging Facility at UC Berkeley. Images were acquired under 543 nm laser illumination focused on the sample using a 20x air objective and collecting 548-685 nm wavelengths using a 54 µm pinhole. Brains were scanned in the z-direction beginning at the top of the brain for 15 planes with 3µm steps. Each image totals the first 45 µm of the brain tissue.

**Table S3.** Optical settings for confocal microscopy

| Dye | Excitation | Emission |
|---|---|---|

| | | |
|---|---|---|
| RhoVR-Halo | 543 nm | 548-685 |

**Confocal imaging analysis.**

Confocal stacks from live prep dye loading of voltage-sensitive dyes were collapsed into a summed z-projections using ImageJ (FIJI). Fluorescence intensity, represented as 8-bit mean pixel values, was measured for the antennal lobe and a region of non-labeled protocerebrum from each brain. The ratio of the AL region's intensity and the nontargeted protocerebrum was then calculated and displayed as fold change above background. No background subtraction was performed in these calculations.

**Carbachol ArcLight functional imaging.**

GH146-Gal4, HaloTag-CD4/CyO; Arclight/Tm2 brains were removed from the animal in AHL-/- and immediately loaded into a perfusion chamber where they were mounted onto a PDL-coated coverslip. Samples were incubated for 3 minutes at room temperature with constant perfusion of AHL+/+ (NaCl 108.0 mM, KCl 5.0 mM, $NaHCO_3$ 4.0 mM, $NaH_2PO_4 \cdot H_2O$ 1.0 mM, Trehalose 2 $H_2O$ 5.0 mM, Sucrose 10.0 mM, HEPES 5.0 mM, $CaCl_2$ 2 $H_2O$ 2.0 mM, $MgCl_2$ 6 $H_2O$ 8.2 mM, perfused at 5 mL/min) before imaging was performed. Perfusion was maintained throughout the experiment. Imaging was performed on an AxioExaminer Z-1 (Zeiss) equipped with a Spectra-X Light engine LED light (Lumencor), controlled with Slidebook (v6, Intelligent Imaging Innovations). Images were acquired with a W-Plan-Apo 20x/1.0 water objective (20x; Zeiss) and focused onto an OrcaFlash4.0 sCMOS camera (sCMOS; Hamamatsu). The acquisition rate of 1 Hz for each experiment with 2 ms exposure times and light power (35 W/cm$^2$) were maintained across all experiments independent of acquisition frequency. A baseline was obtained for one minute, and then brains were stimulated for 30 seconds, followed by a 5-minute recovery period between stimulations.

**Dual Color Functional Imaging.**

GH146-Gal4, HaloTag-CD4; Arclight/Tm2 flies had their cuticle removed as described in the section above and then loaded with 2 µM RhoVR-Halo or 100nM HT-TMR in the presence of 0.2% Pluronic F127 at room temperature for 15 minutes. Following loading, the brains were immediately removed and placed into a perfusion chamber where they were mounted onto PDL-coated coverslips. Samples were incubated for 3 minutes at room temperature with constant perfusion of AHL+/+ (perfused at 5 mL/min) before imaging was performed. Perfusion was maintained throughout the experiment. Imaging was performed on an AxioExaminer Z-1 (Zeiss) equipped with a Spectra-X Light engine LED light (Lumencor), controlled with Slidebook (v6, Intelligent Imaging Innovations). Images were acquired with a W-Plan-Apo 20x/1.0 water objective (20x; Zeiss) and focused onto an OrcaFlash4.0 sCMOS camera (sCMOS; Hamamatsu). The acquisition rate was 1.7 Hz with 125 ms exposure for RhoVR-Halo and 100ms exposure for Arclight. The light power of 13.1 $mW/mm^2$ (RhoVR-Halo) and 19.7 $mW/mm^2$ (Arclight) was maintained across all experiments. A baseline was obtained for one minute and then brains were stimulated in triplicate for 30 seconds followed by a 5-minute recovery period between stimulations.

**Functional imaging data analysis.**

We extracted fluorescence intensity values over time for the antennal lobe using an in-house MATLAB code, and background-subtracted these values. We then corrected the values for bleaching using an asymmetric least squares fit as described previously[6] and added back the baseline value, which was the average of the first 50 frames. Finally, we used the adjusted traces to calculate the % $\Delta F/F_0$, where $F_0$ was defined as the average of frames 2 to 50 from each video. These were then subsequently plotted and displayed using Prism Graph Pad as Mean and Standard Error of the Mean.

**NMJ Imaging and Electrophysiology.**

Third-instar larvae were dissected in ice-cold modified HL3 saline containing (in mM): 70 NaCl, 5 KCl, 10 $MgCl_2$, 10 $NaHCO_3$, 115 mM sucrose, 5 trehalose, and 5 4-(2-hydroxyethyl)-1-piperazineethanesulfonic acid (HEPES) at pH 7.2 as described.[7-8] Guts, trachea, and the ventral nerve cord were removed from the larval body wall. Dissected preparations were then bathed for 15 mins in ice-cold HL3 saline containing 2 µM HT-RhoVR and 0.2% Pluronic F127. The preparation was perfused three times with fresh HL3 saline. For sharp electrode and two-electrode voltage-clamp (TEVC) recordings, electrodes with resistances between 10-35 MΩ were inserted in muscle 6 of segment A2 and A3 in HL3 saline containing 0.4 mM $CaCl_2$. Experiments were conducted using a Zeiss Examiner A1 microscope equipped with a 63x/1.0NA water-dipping objective. Electrophysiological data were acquired using an Axoclamp 900A amplifier, Digidata 1440A acquisition system, and pClamp 10.5 software (Molecular Devices). To elicit EPSPs, an ISO-Flex stimulus isolator (A.M.P.I.) was used to evoke 10 electrical stimulations at 1 Hz with 0.5 msec duration, using stimulus intensities set to avoid multiple EPSPs. To clamp muscles at multiple voltages, the muscle membrane potential was held at -70 mV using a TEVC configuration and clamped from -100 to 0 mV for 500 msec at 10 mV steps. Electrophysiological signals were digitized at 10 kHz and filtered at 1 kHz. Recordings were rejected with input resistances lower than 5 Ωohm or resting potentials more depolarized than -60 mV. Electrophysiology data were analyzed using Clampfit (Molecular Devices). Average holding data for each voltage was calculated for each muscle. Voltage imaging was performed simultaneously with electrophysiological recordings using a PCO sCMOS 4.2 camera at 67 fps (596 x 596 pixels). Voltage imaging was performed using a high-intensity LED (Thor Labs) at 8 individual NMJs during 120-second imaging sessions from two different larvae. Imaging data with severe muscle movements were rejected. Each ROI was manually selected using the outer edge of terminal Ib boutons observed by baseline RhoVR1 signals with ImageJ.[9-10] Fluorescence intensities were measured as the mean intensity of all pixels in each individual ROI. ΔF for an EPSP event or voltage-clamping was calculated by subtracting the baseline RhoVR1 fluorescence level F from the intensity of the RhoVR1 signal during each EPSP event or voltage-clamped at a particular ROI. Baseline RhoVR1 fluorescence of each ROI was defined as average fluorescence in the initial 50 msec of voltage holding or EPSP baseline. ΔF/F was calculated by normalizing ΔF to baseline signal F. The mean intensity at each voltage step was measured as the mean fluorescence intensity of 200 msec during the middle of each voltage step.

**Two-photon excitation spectra and cross-sections.**

The general methods and protocols used for two-photon characterization were published before.[11] Here, we briefly describe them, emphasizing specific details that are different from previously presented methods.

The measurements were done for RhoVR 1 solutions in DPBS buffer pH 7.4 at a concentration of 1 μM.[12] Our experimental setup for two-photon spectral measurements includes a tunable femtosecond laser InSight DeepSee Dual (Spectra-Physics) coupled with a photon-counting spectrofluorimeter PC1 (ISS).[11] The two-photon fluorescence excitation (2PE) spectra were measured by automatically stepping laser wavelength in the range of 700 – 1300 nm and recording total fluorescence intensity at each step in the left emission channel of the PC1 spectrofluorimeter. To eliminate scattered laser light, a combination of short pass filters FF01-770/SP and SP01-633RU-25 (Semrock). The quadratic dependence of fluorescence signal on laser power was observed in this range of wavelengths.

The cross-section $\sigma_{2,A}(\lambda)$ was measured at 840, and 1064 nm using relative method and well characterized reference standards. Rhodamine 6G in methanol was chosen for measurements at 840 nm, with $\sigma_{2,\mathrm{Rh6G}}$ = 43 GM. This values is the average of two independent measurements[13-14] showing good correlation.[11] The measurement at 1064 nm was performed using Rhodamine B in alkaline ethanol as a reference standard. Its cross-section was published relativel to Rhodamine 6G in ethanol, $\sigma_{2,\mathrm{RhB}}/\sigma_{2,\mathrm{Rh6G}}$ = 1.109,[15] 1.000,[16] 1.091,[17] 1.083,[18-19] 1.053,[20] 1.095,[21] with the average $\langle\sigma_{2,\mathrm{RhB}}/\sigma_{2,\mathrm{Rh6G}}\rangle = 1.072 \pm 0.017$. Using this number together with the Rhodamine 6G cross-section in ethanol ($\sigma_{2,\mathrm{Rh6G}}$) obtained after averaging of seven independent measurements: $\langle\sigma_{2,\mathrm{Rh6G}}\rangle = 12.4 \pm 2.6$ GM (see Table 3 of Drobizhev, *et al. Bio-Protocol*, **2020**, *10*, e3498),[11] we calculate $\sigma_{2,\mathrm{RhB}} = 13.3 \pm 2.8$ GM for Rhodamine B in ethanol at 1064 nm and use it as a reference.

For the cross-section measurement, we first collected a total (without monochromator) two-photon excited fluorescence signal $I$ as a function of laser power $P$ for both the sample and reference solutions (samples were held in 3x3 mm cuvettes (Starna) with maximum optical density less than 0.1). The fluorescence was collected at 90° to excitation laser beam through the FF01-770/SP and BLP02-561R-25 (both Semrock) filters, using the left emission channel of a PC1 spectrofluorimeter working in photon counting mode. The power dependences of fluorescence were fit to a quadratic function $I = aP^2$, from which the coefficients $a_\mathrm{S}$ and $a_\mathrm{R}$ were obtained for the sample (index S) and reference (index R) solutions, respectively. Second, the one-photon excited fluorescence signals were measured for the same samples and in the same registration conditions. In this case, a strongly attenuated radiation of a Sapphire 561-50 CW CDRH (Coherent) laser was used for excitation at 561 nm. The fluorescence power dependences for the sample and reference were measured and fit to a linear function: $I = bP$, from which the coefficients $b_S$ and $b_R$ were obtained. The two-photon absorption cross-section was then calculated as follows:

$$\sigma_{2,S}(\lambda_2) = \frac{a_S}{a_R}\frac{b_R}{b_S}\frac{\varepsilon_S(\lambda_1)}{\varepsilon_R(\lambda_1)}\sigma_{2,R}(\lambda_2)$$

Here, $\lambda_1$ is the wavelength used for one-photon excitation (561 nm), $\lambda_2$ is the wavelength used for two-photon excitation (840 or 1064 nm), $\varepsilon_{R,S}(\lambda_1)$ are the corresponding extinction coefficients, measured at $\lambda_1$. This approach allows us to automatically correct for the laser beam properties (pulse duration and spatial intensity distribution), fluorescence collection efficiencies for one- and two-photon modes, PMT spectral sensitivity, differences in quantum yields, and concentrations between S and R solutions.

The extinction coefficients at 561 nm were measured by scaling the corrected fluorescence excitation spectra to the corresponding maximum values: 139,000 $M^{-1}$ $cm^{-1}$ for RhoVR1,[12] 105,000 $M^{-1}$ $cm^{-1}$ for Rhodamine B in alakiline EtOH (measured here), and 100,600 $M^{-1}$ $cm^{-1}$ for Rhodamine 6G in MeOH,[13] and then taking the signal values at 561 nm. Using excitation instead of absorption spectra makes possible to avoid errors due to the contribution of dimers and aggregates to optical density. Finally, the two-photon excitation spectra were scaled to the calculated $\sigma_2$ values with the scaling factor equal to an average scaling factors obtained at two different wavelengths where the $\sigma_2$ was measured.

**Supporting Figures**

**Figure S1.** Immunocytochemistry of HaloTag-CD4 in HEK293T cells.

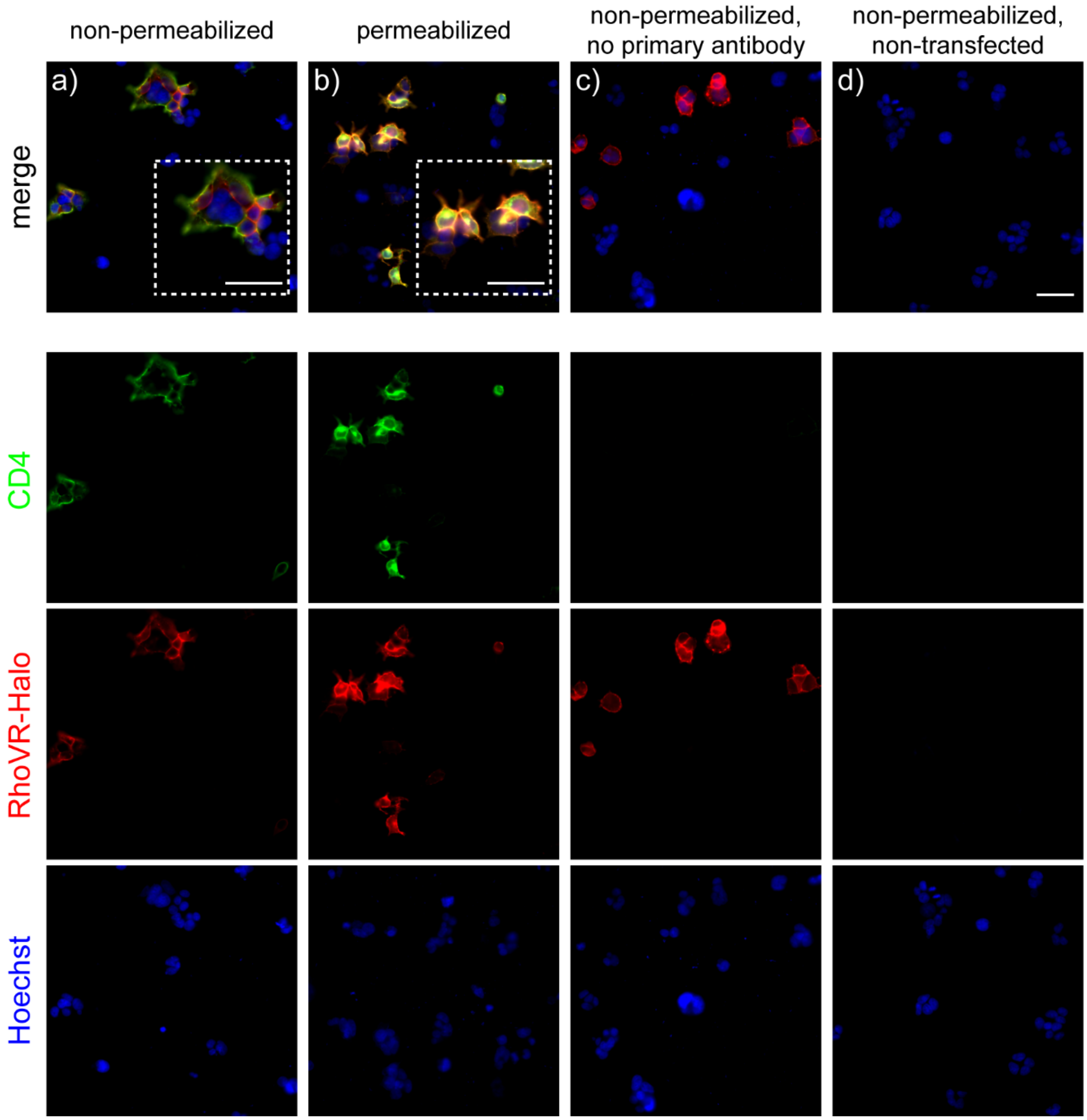

**Figure S1.** Immunocytochemistry of HaloTag-CD4 in HEK293T cells. Widefield epifluorescence images of *post hoc* immunocytochemistry of HEK293T cells expressing HaloTag-CD4, stained with RhoVR-Halo (100 nM) and then treated under **a)** non-permeabilizing conditions (no detergent) or **b)** permeabilizing conditions (0.3% Triton X-100) after fixation. Control cells were treated identically (under non-permeabilizing conditions) and lacked either **c)** primary anti-CD4 antibody or **d)** were not transfected. Green is CD4, red is RhoVR-Halo (100 nM, during live-cell imaging, prior to fixation), and blue is

Hoechst 33342 (at a concentration of 10 μg/mL, equivalent to 16 μM). Scale bar is 50 μm. Insets in panels (**a**) and (**b**) show a zoomed-in region of cells in that panel. Scale bar is also 50 μm.

**Figure S2.** TMR-Halo staining in HEK293T cells.

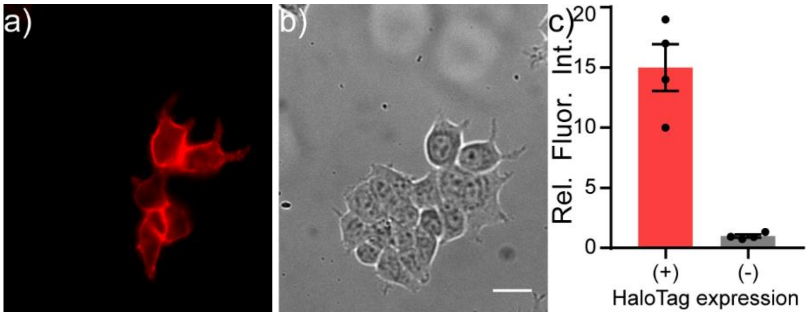

**Figure S2.** TMR-Halo staining in HEK293T cells. Live-cell staining of TMR-Halo in HEK293T cells expressing HaloTag-CD4. Epifluorescence images of HEK293T cells expressing HaloTag-CD4 (CMV promoter) and stained with **a)** TMR-Halo (100 nM, red) and **b)** Hoechst 33342 (1 μM, blue). Scale bar is 20 μm. **c)** Plot of relative fluorescence intensity in cells expressing HaloTag vs. cells that do not express HaloTag. HaloTag-(+) cells were assigned based on a threshold obtained from a non-transfected control. Data are mean ± SEM for n = 4 different coverslips of cells. Data points represent average fluorescence intensities of 40 to 50 cells.

**Figure S3.** Immunocytochemistry in *Drosophila* S2 cells.

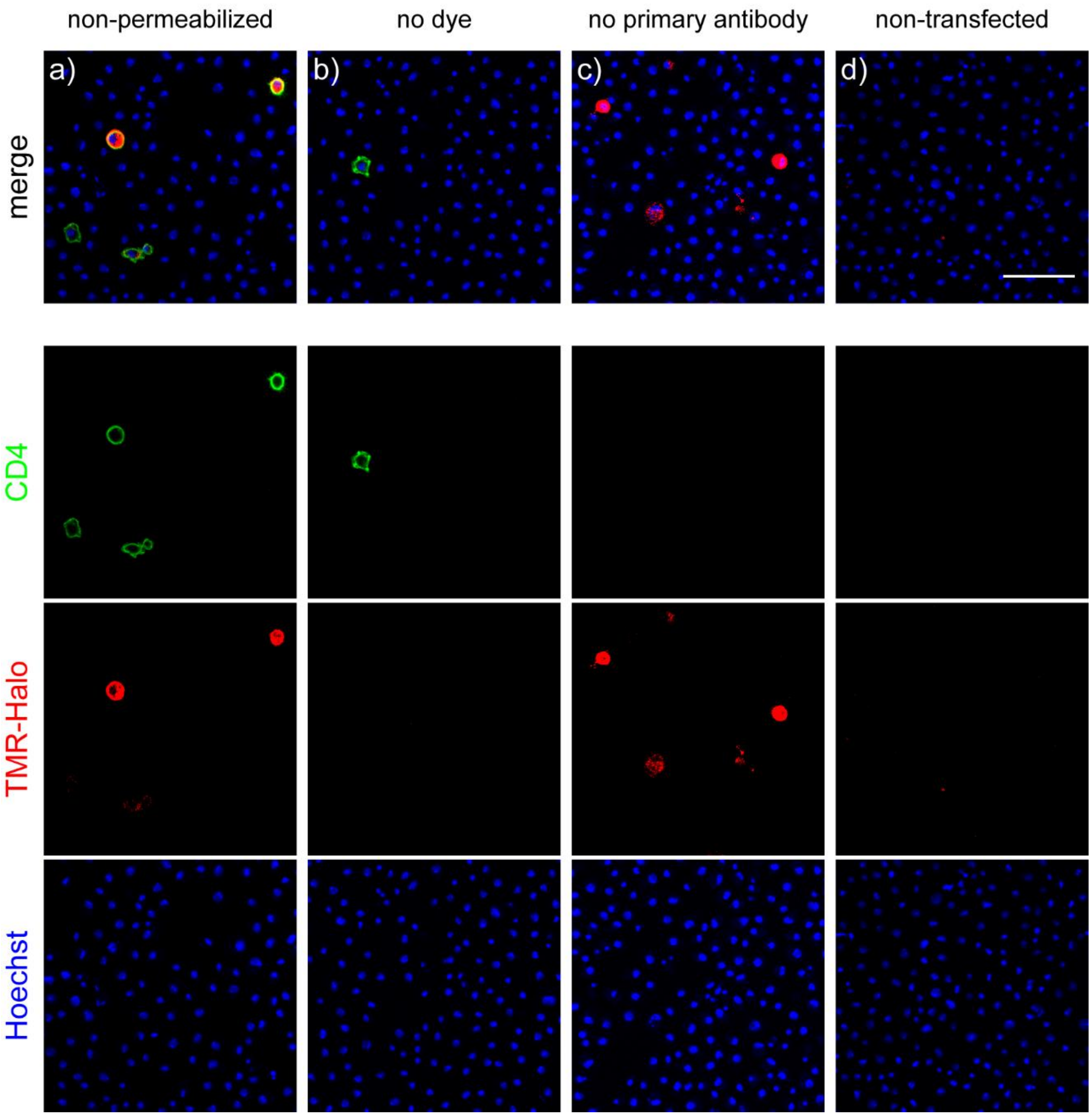

**Figure S3**. Immunocytochemistry in *Drosophila* S2 cells. Confocal images of *post hoc* immunocytochemistry of *Drosophila* S2 cells expressing HaloTag-CD4 (co-transfection with tubP-GAL4 and UAS-HaloTag-CD4 ), stained with TMR-Halo (100 nM, as in **Figure 3** in the main text), and fixed under **a)** non-permeabilizing conditions. Before fixation, control cells were treated either **b)** without TMR-Halo ("no dye"), **c)** with dye, but without primary anti-CD4 antibody, or **d)** with dye, but without transfection. Green is CD4, red is TMR-Halo (100 nM, during live-cell imaging, before fixation), and blue is Hoechst 33342 (at a concentration of 10 μg/mL, is equivalent to 16 μM). Scale bar for all images is 50 μm. Scale bar is also 50 μm.

**Figure S4.** Comparison of voltage sensitivity of RhoVR-Halo with CD4-HaloTag and pDisplay-HaloTag.

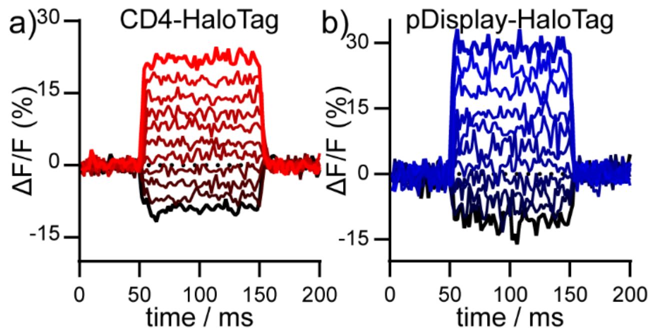

**Figure S4.** Comparison of voltage sensitivity of RhoVR-Halo with CD4-HaloTag and pDisplay-HaloTag. Voltage sensitivity of RhoVR-Halo in HEK293T cells expressing a) HaloTag-CD4 or b) pDisplay-HaloTag. Data are plots of ΔF/F vs. time for a single HEK293T cell expressing either **a)** HaloTag-CD4 (reproduced from Figure 4a in main text) or **b)** HaloTag-pDisplay and stained with RhoVR-Halo. The HEK293T cell was held at -60 mV and then stepped through hyperpolarizing and depolarizing potentials, in 20 mV increments, from -100 mV to +100 mV.

**Figure S5.** Optimization of dye loading in live-fly brains using voltage-sensitive dye RhoVR-Halo.

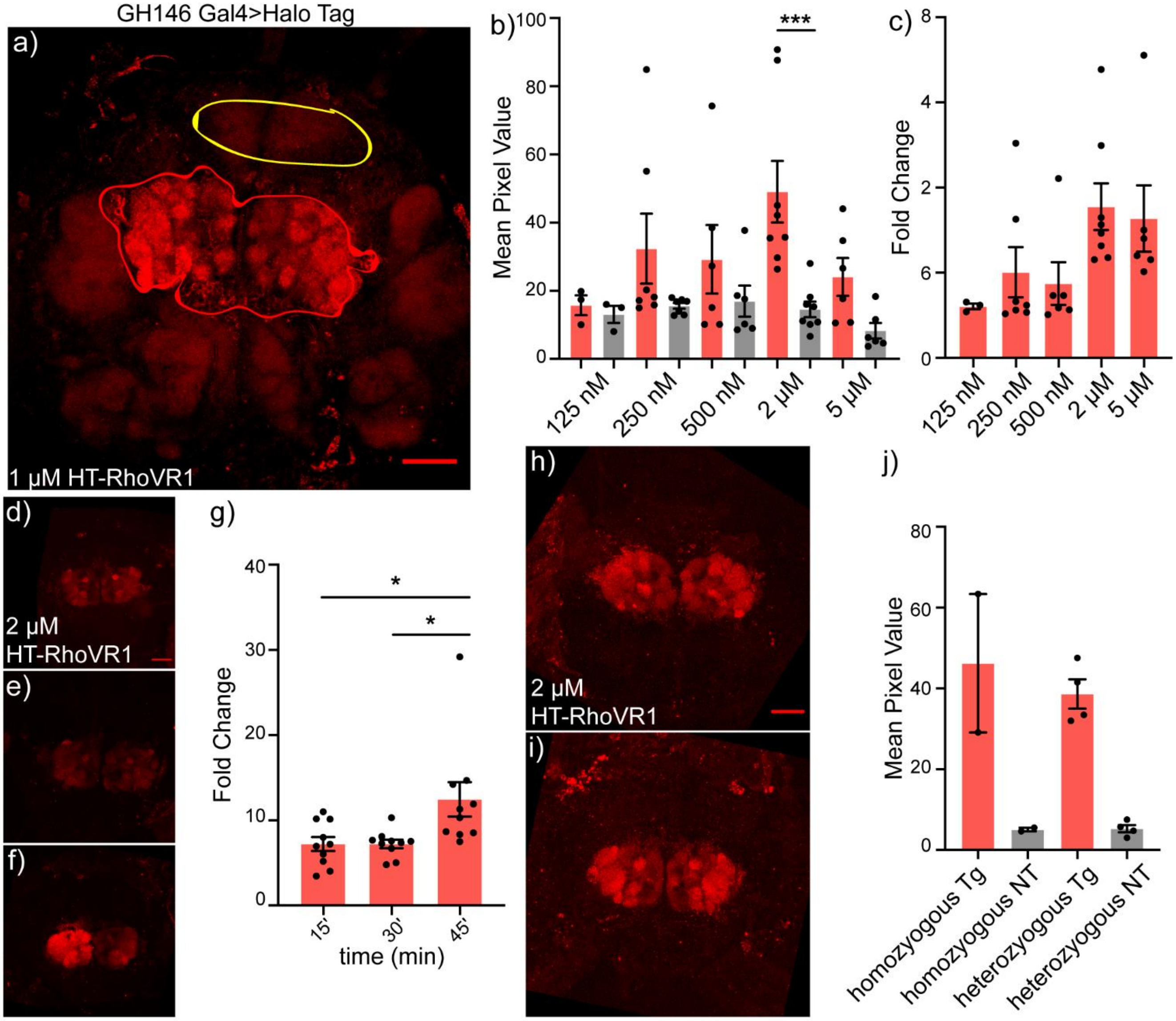

**Figure S5.** Optimization of dye loading in live-fly brains using voltage-sensitive dye RhoVR-Halo. **a)** Maximum confocal z-projection 1 μM RhoVR-Halo loading in GH146 Gal4>HaloTag-CD4 in live explant tissue samples. Regions selected for quantification are circled in red, antennal lobe, and yellow, protocerebrum. Scale is 50 μm. **b)** Quantification of mean fluorescence intensity in GH146 Gal4> HaloTag-CD4 explant brains loaded with varying concentrations of RhoVR-Halo. Red depicts intensities taken from the antennal lobe, while grey depicts intensity taken from the protocerebrum. Data represents at least 6 individual brains per condition (***, $p = 0.0006$). **c)** Quantification of fold change in fluorescence intensity from targeted regions than non-targeted regions in live GH146-Gal4> HaloTag-CD4 loaded with varying concentrations of RhoVR-Halo. Data represent normalized data across at least 6 sample brains per condition. **d)** Maximum z-projection confocal stack of GH146-Gal4> HaloTag-CD4 live explant brain loaded with 2 μM RhoVR-Halo for 15 minutes at room temperature in a live imaging preparation. The scale is 50 μm. **e)** Maximum z-projection confocal stack of GH146-Gal4> HaloTag-CD4 live explant brains loaded with 2 µM RhoVR-Halo for 30 minutes at room temperature in a live imaging preparation. The scale is the same as d. **f)** Maximum z-projection confocal stack of GH146-Gal4> HaloTag-CD4 live explant brains loaded with 2 µM RhoVR-Halo for 45 minutes at room temperature in a live imaging preparation. The scale is the same as d. **g)** Quantification of fold fluorescence intensity change in GH146-Gal4>HaloTag-CD4 treated with 2 µM RhoVR-Halo for 15,30 or 45 minutes. Data represents normalized fluorescence intensity across at least 9 independent samples per condition. **h)** Maximum z projection of a confocal stack from RhoVR-Halo loaded GH146-Gal4> HaloTag-CD4 heterozygote live explant brains. The scale is 50 µm. **i)** Maximum z projection of a confocal stack from RhoVR-Halo loaded GH146-Gal4> HaloTag-CD4 homozygote live explant brains. The scale is the same as h. **j)** Quantification of fold intensity change in RhoVR-Halo loaded GH146-Gal4> HaloTag-CD4 heterozygotes and homozygotes.

**Figure S6.** Imaging carbachol-induced depolarization in *Drosophila* projection neurons in live explants using ArcLight.

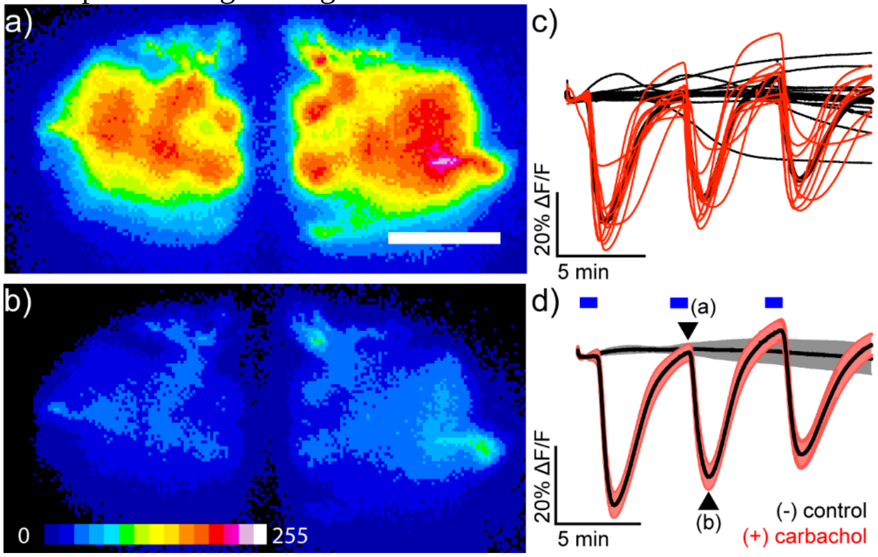

**Figure S6.** Imaging carbachol-induced depolarization in *Drosophila* projection neurons in live explants using ArcLight. Epifluorescence images of live explant *Drosophila* brain expressing ArcLight in antennal lobe projection neurons (GH146-GAL4,HaloTag-CD4/CyO>ArcLight/TM2) **a)** immediately before and **b)** 30 s after stimulation with 100 μM carbachol. Scale bar is 50 μm. Image is pseudo-colored, and the scale bar indicates 8-bit pixel grey values. **c)** Plot of ArcLight fluorescence (ΔF/F) vs. time for individual fly brains in response to three 100 μM carbachol stimulations (red, n = 10) and control (vehicle only) stimulations (black, n = 11). Each trace represents one individual brain. **d)** Plot of mean ArcLight fluorescence (ΔF/F). Data are mean ± SEM. Blue bars represent the addition of vehicle (control) or carbachol.

**Figure S7.** Imaging osmolarity-induced hyperpolarizations in ISNs in *Drosophila* using Arclight voltage indicator.

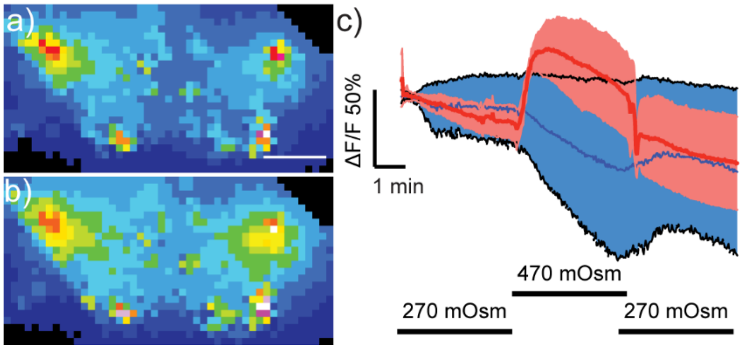

**Figure S7.** Imaging osmolarity-induced hyperpolarizations in ISNs in *Drosophila* using Arclight voltage indicator. Spinning disk confocal maximum z-projections of live explant *Drosophila* brain expressing Arclight in the ISNs (Nanchung-Gal4, UAS-HaloTag/ UAS-Arclight; TM2/TM6B) **a)** before and **b)** after stimulation with high osmolarity hemolymph (470 mOsm). Scale bar is 50 µm. Image is pseudo-colored, and the scale bar indicates 8-bit pixel grey values. **c)** Plot mean of fluorescence (%ΔF/F) vs. time in response to either a high osmolarity stimulation (red, n=8) or vehicle control (blue, n=8). Data are mean ± SEM Black bars below indicate the stimulation time course switching from 270 mOsm to 470 mOsm.

**Figure S8.** Imaging osmolarity-induced hyperpolarizations in ISNs in heterozygous transgenic *Drosophila* using RhoVR-Halo voltage indicator.

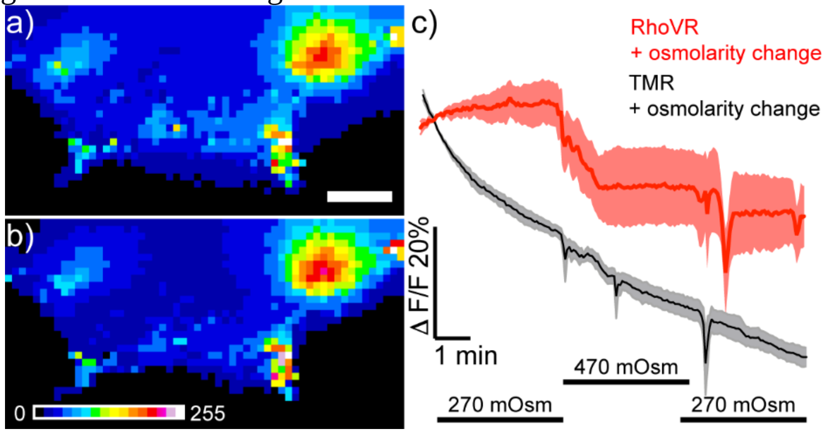

**Figure S8.** Imaging osmolarity-induced hyperpolarizations in ISNs in heterozygous transgenic *Drosophila* using RhoVR-Halo voltage indicator. Spinning disk confocal maximum z-projections of live explant *Drosophila* brain expressing HaloTag-CD4 in the ISNs (Nanchung-Gal4, UAS-HaloTag/CyO; TM2/TM6B) **a)** before and **b)** after stimulation with high osmolarity hemolymph. Scale bar is 50 µm. Image is pseudo-colored, and the scale bar indicates 8-bit pixel grey values. **c)** Plot mean of fluorescence (%ΔF/F) vs. time in response to one osmolarity simulation of either RhoVR-Halo (red, n = 5) or HT-TMR (black, n=5) loaded brains. Data are mean ± SEM Black bars below indicate the stimulation time course switching from 270 mOsm to 470 mOsm.

**Figure S9.** Two-photon spectra of RhoVR 1.

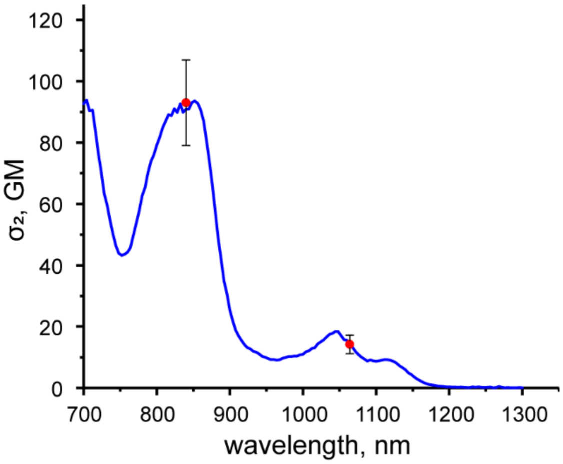

**Figure S9.** Two-photon absorption spectra of RhoVR 1. Two-photon excitation spectrum (solid blue line) is scaled to the absolute two-photon absorption cross-section values measured at selected wavelengths (red dots) ($\sigma_2$, in units of Göppert-Mayer, 1 GM = $10^{-50}$ $cm^4$ s $molecule^{-1}$). Error bars are associated with the absolute absorption cross-section value. A machine-readable file (.csv) is included in the supplementary files.
